# Novel insights into potential therapeutic targets and biomarkers using integrated multi-*omics* approaches for dilated and ischemic cardiomyopathies

**DOI:** 10.1101/2020.12.15.422946

**Authors:** Austė Kanapeckaitė, Neringa Burokienė

## Abstract

At present heart failure treatment targets symptoms based on the left ventricle dysfunction severity; however, lack of systemic studies and available biological data to uncover heterogeneous underlying mechanisms on the scale of genomic, transcriptional and expressed protein level signifies the need to shift the analytical paradigm toward network centric and data mining approaches. This study, for the first time, aimed to investigate how bulk and single cell RNA-sequencing as well as the proteomics analysis of the human heart tissue can be integrated to uncover heart failure specific networks and potential therapeutic targets or biomarkers. Furthermore, it was demonstrated that transcriptomics data in combination with minded data from public databases can be used to elucidate specific gene expression profiles. This was achieved using machine learning algorithms to predict the likelihood of the therapeutic target or biomarker tractability based on a novel scoring system also introduced in this study. The described methodology could be very useful for the target selection and evaluation during the pre-clinical therapeutics development stage. Finally, the present study shed new light into the complex etiology of the heart failure differentiating between subtle changes in dilated and ischemic cardiomyopathy on the single cell, proteome and whole transcriptome level.

**HIGHLIGHTS:**

- First report of an integrated multi-omics analysis for dilated and ischemic cardiomyopathies.
- Identification of metabolic and regulatory network differences for the two types of cardiomyopathies.
- Introduction of a new scoring system to evaluate genes based on the size of their network and disease association.
- Two-step machine learning pipeline to uncover potential therapeutic target clusters.

## Introduction

Cardiovascular disease (CVD) is the leading cause of death globally; however, both investment and efforts in CVD drug development are declining. This contrasts sharply with funding and drug approvals for other indications, such as oncology^1,2^. While there are many factors contributing to this trend^3^, low tolerance for side-effects and lack of good biomarkers are some of the key challenges in implementing new therapies. Thus, all of this calls to revisit currently used approaches in the therapy development for CVD. Specifically, combining high-throughput RNA-sequencing (RNA-seq), proteome analysis and biological data mining could potentially facilitate the identification of new therapeutic targets by deconvoluting complex pathways involved in the pathological processes. Subsequently, gaining a better understanding of the disease etiology on the molecular level could also be advantageous for a better monitoring of the pathology progress and treatment efficacy.

Heart failure (HF) affects approximately 40 million people globally as recorded in 2015 and estimated 2% of adult population is suffering from HF^4^. HF dominates in the elderly population with the incidence rate being 6-10% for those over 65 years old and more than 10% for the population older than 75^4,5^ with men showing a higher predisposition for CVD^6^.

Most cardiomyopathies have complicated underlying causes where chronic or poorly controlled hypertension can lead to increased afterload resulting in higher cardiac workload which in turn can precipitate hypertrophy of the left ventricle. Decreased heart contractility and output in CVD can also be caused by a direct ischemic damage to the myocardium which induces further scar formation and tissue remodelling^7^. Hypertension, ischemic cardiomyopathy (IC) and dilated cardiomyopathy (DC) precedes later-stage heart failure with reduced ejection (HFrEF)^8,9^. HFrEF encompasses a diverse pathologic spectrum and is a good case example for when long-standing paradigms of a single common pathway^9,10^ do not provide an adequate measure of the pathology development or progression. That is, most current therapies for HFrEF do not specifically focus on disease etiology or in-depth differentiation^9^; thus, the heterogenous nature of HF remains insufficiently addressed. As a result, the need of new therapeutic insights and an improved analysis of underlying HF mechanisms were the divining forces behind this study to develop a novel approach with integrated multi-*omics* and machine learning methods.

The dramatic expansion of RNA-seq and metabolomics screening capabilities provides an excellent resource for an in-depth look into cardiomyopathies. Moreover, while cardiac sample collection cannot always be optimal and there are technical variations, a robust growth in novel statistical approaches allows researchers to better glean information from noisy datasets and clean the data from technical errors or batch effects. To demonstrate how multi-*omics* approaches can help to uncover the intricate biological mechanisms of pathological processes, the human left ventricle was selected as a case study since current HF treatment relies on targeting the associated symptoms with left ventricle failure without taking into account the heterogeneity of underlying mechanisms^2,9,11,12^. Thus, the central hypothesis was that the analysis of transcriptome and proteome data of human heart tissue with novel statistical and machine learning methods would help to profile potential therapeutic targets or affected pathways (Fig. 1).

**Figure 1.**
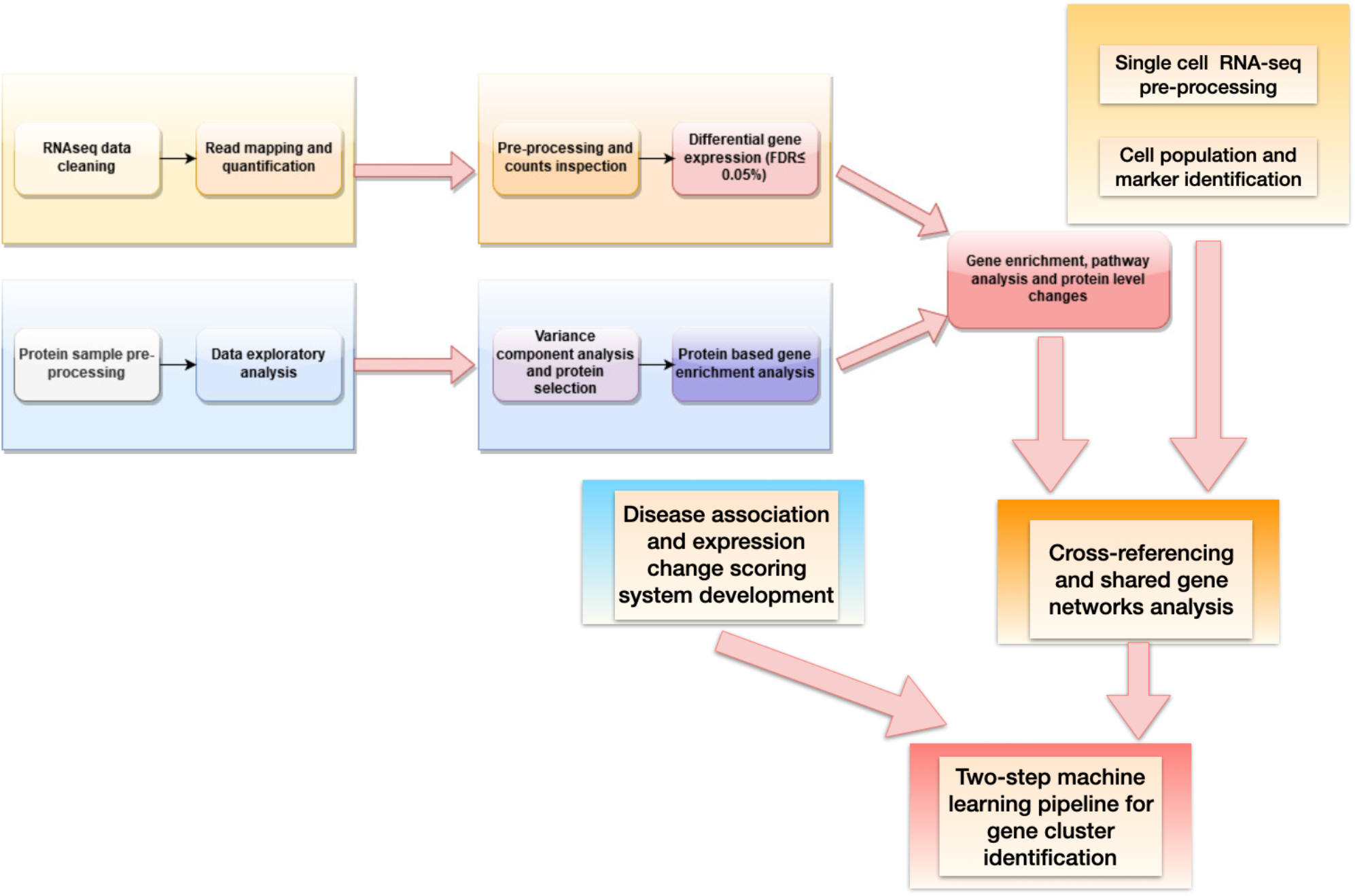
Diagram showing the steps for data processing and integration.

Relatively small sample was used in this study to better simulate what happens during clinical trials and studies focusing on patients; that is, it was necessary to replicate the situation where the number of samples might be limited, show sex-dependent variation and not perfect age group matching. Thus, twelve random human left ventricular RNA-seq samples were selected to form three categories: non-failing (Healthy), dilated (DC), and ischemic (IC), with approximately equal representation for all ages and sexes; similar sets of samples were selected for proteome analysis (Supplementary Tables 1&2). This was followed by a multi-analytical approach to filter and probe differentially expressed genes and generated datasets were used for gene enrichment and pathway analysis as well as clustering using two-step machine learning approaches (Fig. 1).

## Methods

### Sample selection

Publicly available datasets were used to randomly select twelve human left ventricular RNA-seq samples (PRJNA477855^9,13^, EBI: European Nucleotide Archive^14^) which were categorised to form: non-failing (Healthy), dilated (DC), and ischemic (IC) cardiomyopathy groups; similar sets of samples were selected for the proteome analysis (PXD008934^15^, EMBL-EBI: PRIDE^16,17^) (Supplementary Tables 1&2) with matched representation for all ages and sexes. Single cell RNA-seq of the murine non-myocyte cardiac cellulome (E-MTAB-6173^18^) was downloaded from ArrayExpress database^19^. Human left ventricular myocardium was downloaded from publicly available Visium data from 10X Genomics^20^.

### RNA-seq data pre-processing and exploratory analysis

The - number of reads per sample ranged from approximately 40 to 65 million averaging to 59 million. Reads were filtered for quality and trimmed using Trimmomatic tool^21^ and aligned to the human genome reference GRchr37/hg19^22^ using HISAT2^23^ with 95% average alignment rate. Ensembl GRCH37/ hg19^22^ GTF was used for featureCounts^24^ tool to count reads based on genomic features. Quality control was performed both pre-and-post-alignment using MultiQC^25,26^ and Integrative Genomics Viewer (IGV)^27^ was used to additionally inspect alignments. Single cell counts were acquired after the raw data was processed with Cell Ranger version 1.3 (10x Genomics)^28^.

### Differential expression analysis

R Studio 3.6^29^ environment was used for raw RNA-seq counts pre-processing and further analysis using package DESeq2^30,31^ as well as dependent packages for graphical processing and data manipulation. Raw count variability (Supplemetary Fig. 2) was evaluated using statistical analysis for outliers (count density distribution analysis, distance clustering, Bland–Altman (MA) plot analysis) and lowly expressed genes were removed from the samples to better control for the false discovery rate (p-adjusted, FDR; Benjamin-Hochberg correction)^30,31^. A linear fitting model was used to compare different sample groups based on the disease status while controlling for gender differences. For other downstream analyses, for example, sample distribution visualisation, the regularised logarithm (rlog) was used to adjust for the sample dependence of the variance on the mean. Seurat R package^32^ was used to analyse single cell data maintaining mitochondrial DNA content at <5% for non-cardiomyocyte samples and <40% for cardiomyocites, all appropriate control steps (unique counts, count distribution per cell and per gene) were used to inform the downstream analyses. R packages: SingleR^33,34^, CellDex^35^ and Clustermole^36^ were used to determine cell types for the cell clusters in the single cell sequencing datasets. Where available the following contrasts, dilated cardiomyopathy vs healthy (DC vs Healthy) and ischemic cardiomyopathy vs healthy (IC vs Healthy), were analysed to uncover specific changes for a disease when the observed changes are weighed against normal samples.

**Figure 2.**
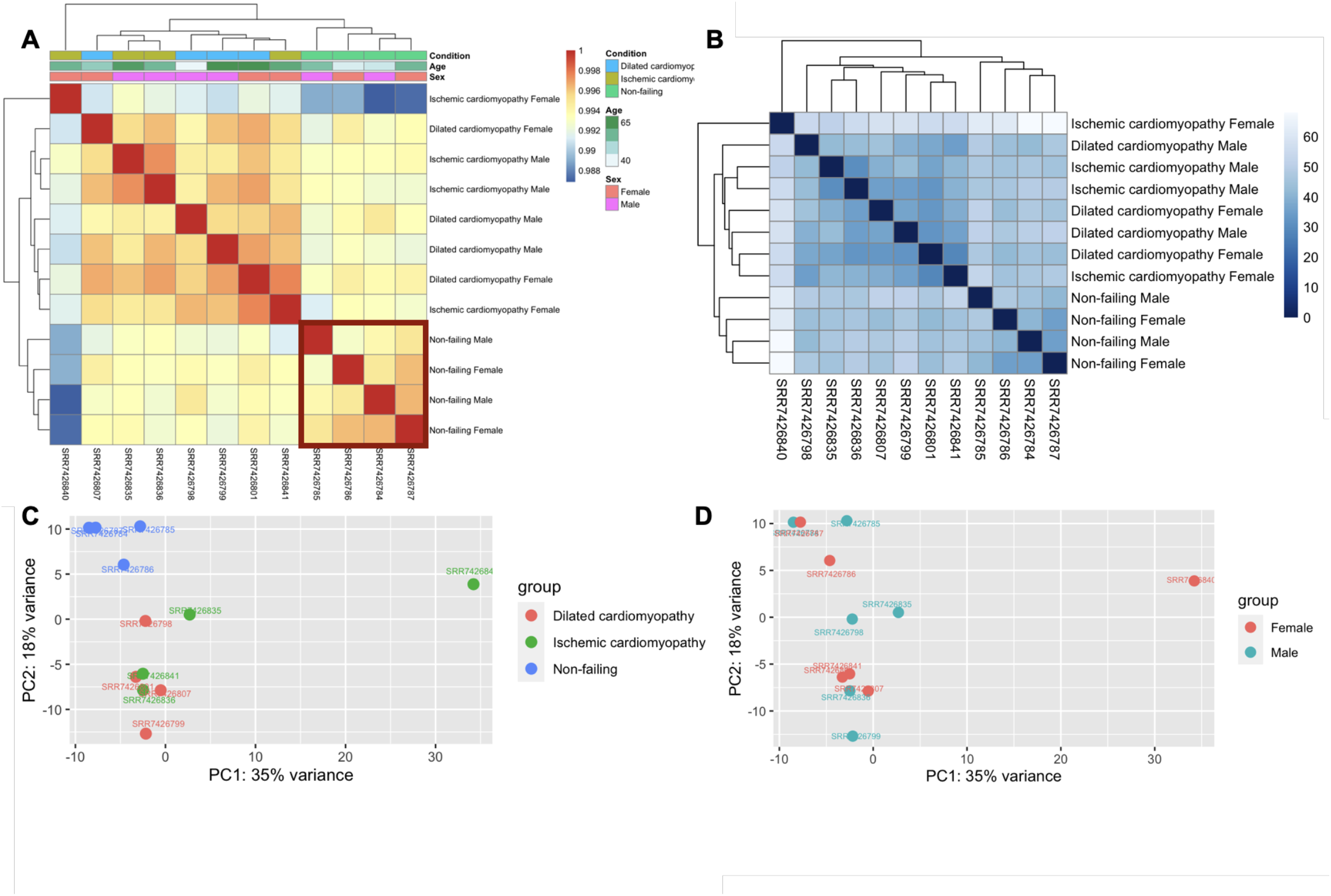
Human left ventricle bulk RNA-seq (PRJNA477855) gene count clustering and distribution analysis showing Spearman correlation calculated distances (A) and euclidean distances (B) for rlog transformed counts using complete-linkage hierarchical clustering method; sample distributions across top two principal components are shown in the PCA plot grouping by condition (C) and gender (D).

### Protein level analysis

Protein abundance data was retrieved from earlier raw spectra analyses using MaxQuant version 1.5.3.30^37^ integrated with the Uniprot human database^38^ search. Label-free quantification (LFQ) intensity values were used in lieu of protein abundance and were pre-processed to remove proteins with median distributions across all samples that were equal to 0 LFQ. LFQs were scaled by a factor of 10^−6^ prior to DESeq2 based normalisation and model fitting to find differences between conditions while controlling for gender effects.

### Gene enrichment and pathway modelling

R annotation libraries: AnnotationDbi^39^, ClusterProfiler^40,41^ as well as DEGReport^42^ and dependent packages were used for gene ontology and pathway analysis. Specifically, a hypergeometric test for gene set enrichment analysis by mining directed acyclic association graphs was performed to select enriched gene sets and associated pathways. Reactome^43^, Open Targets^44^, STRING (version 11, score_threshold=200)^45^ and STITCH^46^ databases were used for data mining to build interactor networks.

### Machine learning and disease centric scoring

For the initial clustering Gaussian mixture models (GMMs) were chosen since they function as a density estimator to establish cluster patterns. The probabilistic nature of GMM was best suited to perform parameter separation^47,48^. Identified clusters with GMM where isolated and subjected to agglomerative hierarchical clustering^49,50^ (implemented via Hclust R functionality) since this method is the most optimal to find small sub-clusters. GMMs (with the following parameters: max_iter=1000, covariance_type=‘full’ or ‘spherical’, tol=0.001, random_state=0) were implemented to cluster genes based on their scaled Log2 fold change (LFC_Score_ >|1.5|) and the number of interactors45 (1 eq.). Scaling factor was determined by the cumulative score of multiple mined resources (Open Targets^44^) where a gene was assigned a value (from 0 to 1) based on its probabilistic links to a specific disease. The number of interactors was identified using STRING database of known protein-protein interactions^45^. GMM clustering evaluation was performed using probabilistic statistical measures quantifying the model performance for the different number of clusters. Evaluation parameters were based on Akaike information criterion (AIC)^51^ and the Bayesian information criterion (BIC)^51^. AIC allows to estimate the in-sample error as well as information loss, while BIC addresses the overfitting based on the likelihood function.

The number of components for clustering and correction of the over-fitting was established based on using AIC and BIC information criterion values. Python Scikit-Learn GMM (scikit-learn 0.22.2)^52^ was used to determine Gaussian mixture modelled density and distribution of selected gene parameters. Prior to clustering for a selected contrast, scored genes were pre-filtered to remove minimally changed values (LFC_Score_ >|1.5|). Following this, selected GMM clusters were subjected to agglomerative hierarchical clustering and the hierarchical clustering dendograms were divided into sub-groups based on the optimal number determined by Silhouette and Elbow methods^53,54^.

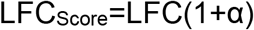

**Equation 1.** Log2 Fold Change_Score_ equation defines a scaled LFC (log2 fold change) value for a given contrast where α is a value showing the strength of disease association for a given gene.

### Graphs and statistical analyses

R Studio 3.6^29^ was used for all plots and data processing. Python Scikit-Learn (scikit-learn 0.22.2)^52^ was used for GMM implementation with density plot visualisation.

## Results

### Bulk RNA-seq captured disease specific gene changes in dilated and ischemic heart conditions

Exploratory analysis of the human left ventricle bulk RNA-seq data (PRJNA477855^13^) revealed that the sample count distribution and coverage depths were consistent (Supplementary Fig. 1&2) without any marked batch effects. However, clustering analysis (Fig. 2, A&B) indicated that samples were relatively homogenous based on their gene expression with only Non-failing (healthy) group showing the clearest separation. Moreover, dilated and ischemic cardiomyopathy groups were intertwined without minimal subdivision. This trend was also reflected in the principal component analysis (PCA) (Fig. 2, C) where the disease groups not only had a marked overlap but also the intra-sample variability was higher compared to the healthy group.

Despite high homology between samples, the observed differences for various contrast groups, e.g. disease state vs healthy, can still reveal genes involved in biologically meaningful processes if they had a marked upregulation or downregulation. This assumption was confirmed by the proportion of significantly changed genes for dilated (11.1%) and ischemic (17.6%) pathological states when contrasted to the healthy tissue (Fig. 3).

**Figure 3.**
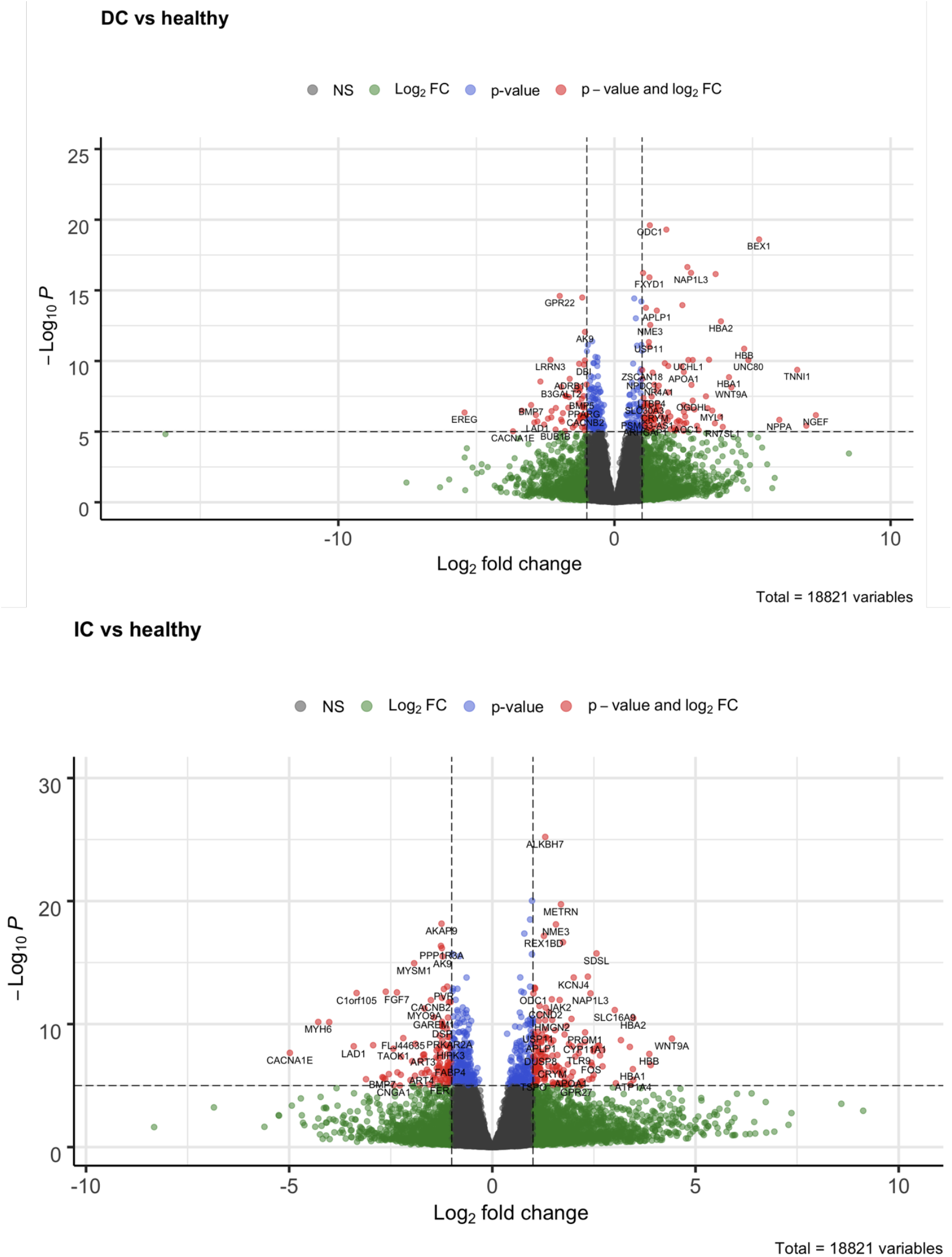
Human left ventricle bulk RNA-seq (PRJNA477855) significantly changed gene count Volcano plots where IC – ischemic cardiomyopathy and DC – dilated cardiomyopathy, FDR p-adjusted<0.00001 and Log fold change (LFC)>|2|.

Interestingly, the genes that changed significantly for each investigated contrast showed some overlap which can be attributed to the complex nature of the regulatory pathways involved as some gene networks inadvertently become shared between the pathological processes^6,11,55^. Both unique and full sets of significantly changed genes per contrast group (Supplementary Fig. 4) were used to enrich for marker genes as it was necessary to examine shared and disease specific expression patterns in the pathology. Genes that had the most notable change based on p-adjusted value when comparing dilated cardiomyopathy vs healthy samples showed a clear separation for these two conditions (Fig. 4, A). However, the same set of genes did not show such a pattern in ischemic disease. When selecting genes based on the lowest p-adjusted value for the contrast of ischemic cardiomyopathy vs healthy heart samples (Fig. 4, B), the ischemic heart sample genes formed a separate cluster while healthy and dilated cardiomyopathy samples were dispersed and relatively similar in their expression values. Further investigation of the most significantly changed genes in dilated cardiomyopathy (Fig. 4, A) revealed that Ribosomal Protein S17 (RPS17) expression is the most notably changed. While ribosomal proteins might be a left-over due to the sample preparation, there is emerging evidence of ribosomal protein expression and/or mutational changes being involved in numerous diseases; RPS17 and other ribosomal protein family members were linked to tissue morphogenesis and structural changes^56^. Since there was no other over-representation for ribosomal genes, it is possible that the observed expression levels might be biologically meaningful. Other groups of genes, such as SLIT And NTRK Like Family Member 4 (SLITRK4) and Glycosyltransferase 8 Domain Containing 2 (GLT8D2), have been reported to have links to tissue structural changes^57^. Upregulated Myozenin-1 (MYOZ1), Enolase 2 (ENO2) and Bone Morphogenetic Protein 2 (BMP2) are all linked to heart tissue hypertrophy or were identified as potential biomarkers in the disease^44,58^. These findings are especially interesting when compared to the downregulated genes, specifically Carbonic Anhydrase 11 (CA11), Intercellular Adhesion Molecule 3 (ICAM3) and ELOVL Fatty Acid Elongase 2 (ELOVL2), as these molecules have been associated with heart failure, vascular injury and changes in tissue metabolism^44,59–61^.

**Figure 4.**
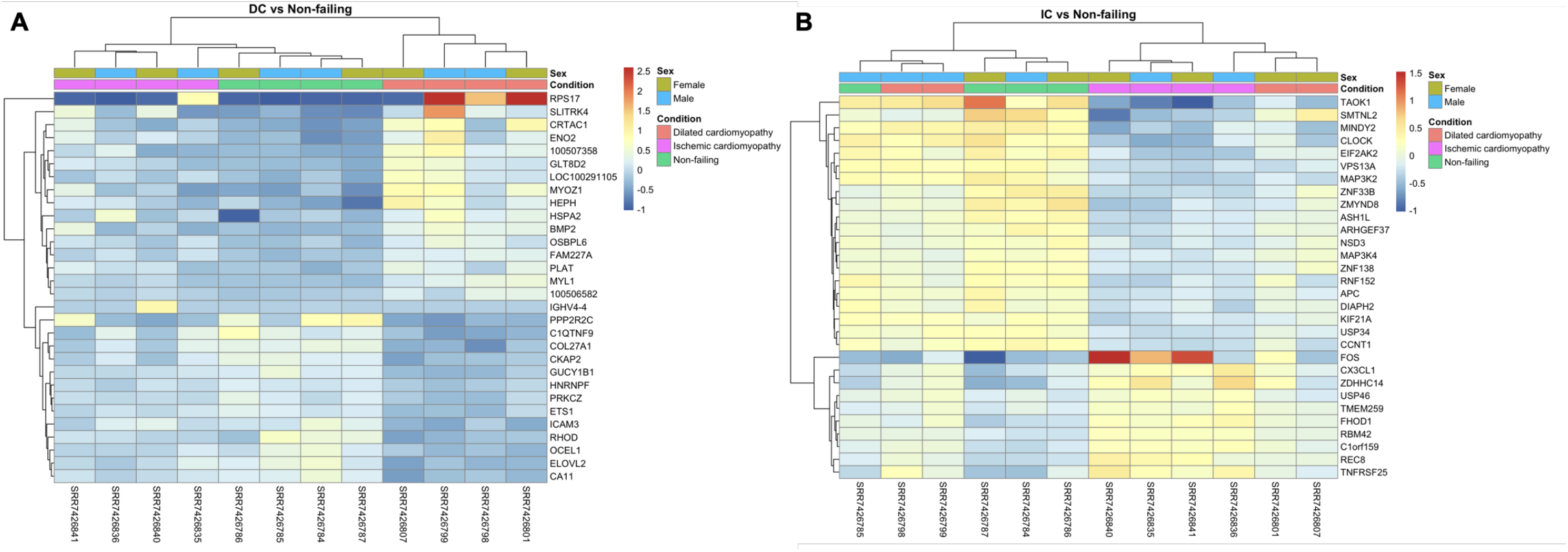
Heatmap for significantly changed genes that are unique for dilated cardiomyopathy (DC) vs healthy samples (A) and ischemic cardiomyopathy (IC) vs healthy samples (B) where values are shown for all conditions. Counts reported are rlog transformed and mean standardised per gene.

In contrast to the dilation of the heart, ischemic conditions were found to be dominated by immune system, fibrosis and cell proliferation linked genes, namely C-X3-C Motif Chemokine Ligand 1(CX3CL1), Proto-oncogene c-Fos (FOS), Transmembrane Protein 259 (TMEM259), REC8 Meiotic Recombination Protein (REC8) and Formin Homology 2 Domain Containing 1 (FHOD1), that were significantly expressed and some of the genes, such as CX3CL1 and TMEM259, are candidate genes for novel biomarkers and/or therapeutic targets for the ischemic heart disease^17,62–64^. The group of downregulated genes in ischemia, for example, TAO Kinase 1 (TAOK1) and MINDY2 (Lysine 48 Deubiquitinase 2), are categorised as involved in some inflammatory processes^65–73^.

These observations reflect not only the complex biological mechanisms underlying the heart disease but also highlight that target gene selections have to be made based on multiple criteria: significance, observed differences and uniqueness to a disease phenotype.

Exploring uniquely and significantly changed genes in dilated or ischemic cardiomyopathy but ranking based on the fold change (Supplementary Figure 5), we can immediately see that dilated cardiomyopathy showed interesting metabolic patterns, such as the upregulation of 5-HT transporter (serotonin transporter, SLC6A4) with dependence on sodium- and chloride-movement across the membrane as well as an increase in Cytochrome P450 Family 3 Subfamily A Member 5 (CYP3A5) expression; RPS17 also belonged to this LFC ranked category. As in previous p-adjusted value dependent categorisation, the ischemic heart tissue had a more pronounced signature of immune process involvement, for example, Major Histocompatibility Complex, Class I, C (HLA-C) and Immunoglobulin Lambda Variable 6-57 (IGLV6-57) (Supplementary Figure 5). It was also further demonstrated that the significantly changed genes for the contrasts of interest showed no marked sex biases (Supplementary Fig. 6&7); thus the following analyses focused on the biological processes driving the observed changes in the expression patterns.

### Human heart left ventricle bulk RNA-seq revealed clear pathological process bifurcation for dilated and ischemic cardiomyopathies

Emerging differences between the iscemic and dilated heart were further cemented by exploring gene enrichment and associated biological processes. Not surprisingly enriched processes for the dilated heart (Fig. 5, A&B) belonged to myocardium remodelling, ventricular cardiac muscle tissue morphogenesis and muscle tissue development. However, a specific set of enriched process was found for the genes that were only significantly changed in dilated and not ischemic heart (Fig. 5, C&D), these genes are involved in microtubule, myofibril, sarcromere and contractile fibre processes.

**Figure 5.**
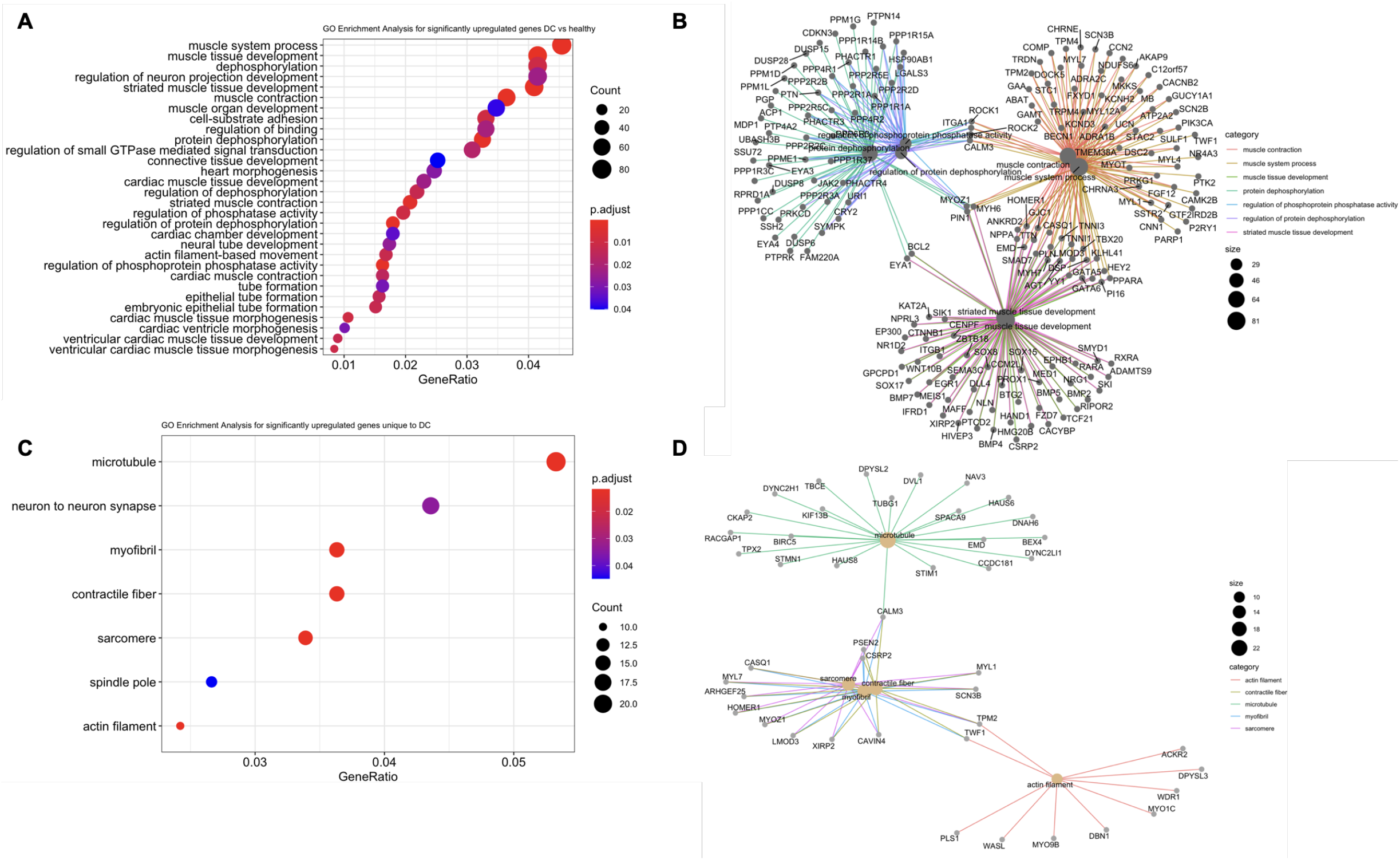
Enrichment analysis for all significantly changed genes in the DC (dilated cardiomyopathy) vs Healthy contrast group where graph A shows enriched cellular processes and map B provides visualisation of the top highest ranking processes and corresponding genes. Gene set size that was enriched and enrichment p-adjusted value provided with the plots. Panels B and C represent gene enrichment for unique and significantly changed genes in the DC vs Healthy group.

There are 64 genes (Supplementary Table 3) that were not only significantly changed when comparing dilated heart state with a healthy sample but also clustered into distinct cellular processes (Fig. 5, C&D). Some of those genes, namely Myosin light chain 1 (MYL1), Dynein axonemal heavy chain 6 (DNAH6), MYOZ1 and Atypical chemokine receptor 2 (ACKR2), showed a significant upregulation in a disease state and could be of interest as potential therapeutic targets or biomarkers^44^.

Enrichment of the gene networks for ischemic conditions revealed a specific involvement in heart ventricular cardiac muscle tissue morphogenesis and broader metabolic functions, such as GTPase activity linked processes (Fig. 6, A&B). While tissue remodelling is expectedly shared between ischemic and dilated cardiomyopathy, there were more subtle differences in ischemic conditions that hint toward ER stress and inflammatory processes (Fig. 6, C&D). For example, Spingomyelinase SMPD3 has been previously implicated in Golgi vesicular protein transport where the inactivation of this enzyme disrupted proteostasis leading to ER stress^59,74–76^. At the intersection of ER stress and immunological processes there was another significantly upregulated gene, Formyl peptide receptor 2 (FPR2), which downregulation has been shown to alleviate oxidative and inflammatory burden^77,78^ (Supplementary table 4). Intriguingly, there was a number of chemokine ligands (e.g., CXCL11, CXCL10 and CCL5) that were highly expressed as well as some chemokine receptors (e.g., CXCR3 and CCR7) and other markers, such as CD2 (Supplementary Table 4). While chemokine ligands can be expressed on a number of cells^79^, the receptor role is more associated with T-cells and other lymphoid cells or tissues^80–83^. CD2 marker expression is very clearly ascribed to T-cells and complex immune regulatory environment^80,84,85^, these findings likely point to a heterogenous nature of the heart samples with other lymphoid cells infiltrating affected tissues. Nevertheless, there is a clear shift in the ischemic tissue state with an increased inflammatory burden and multiple regulatory mechanisms engaged (e.g., CX3CL1)^86^.

**Figure 6.**
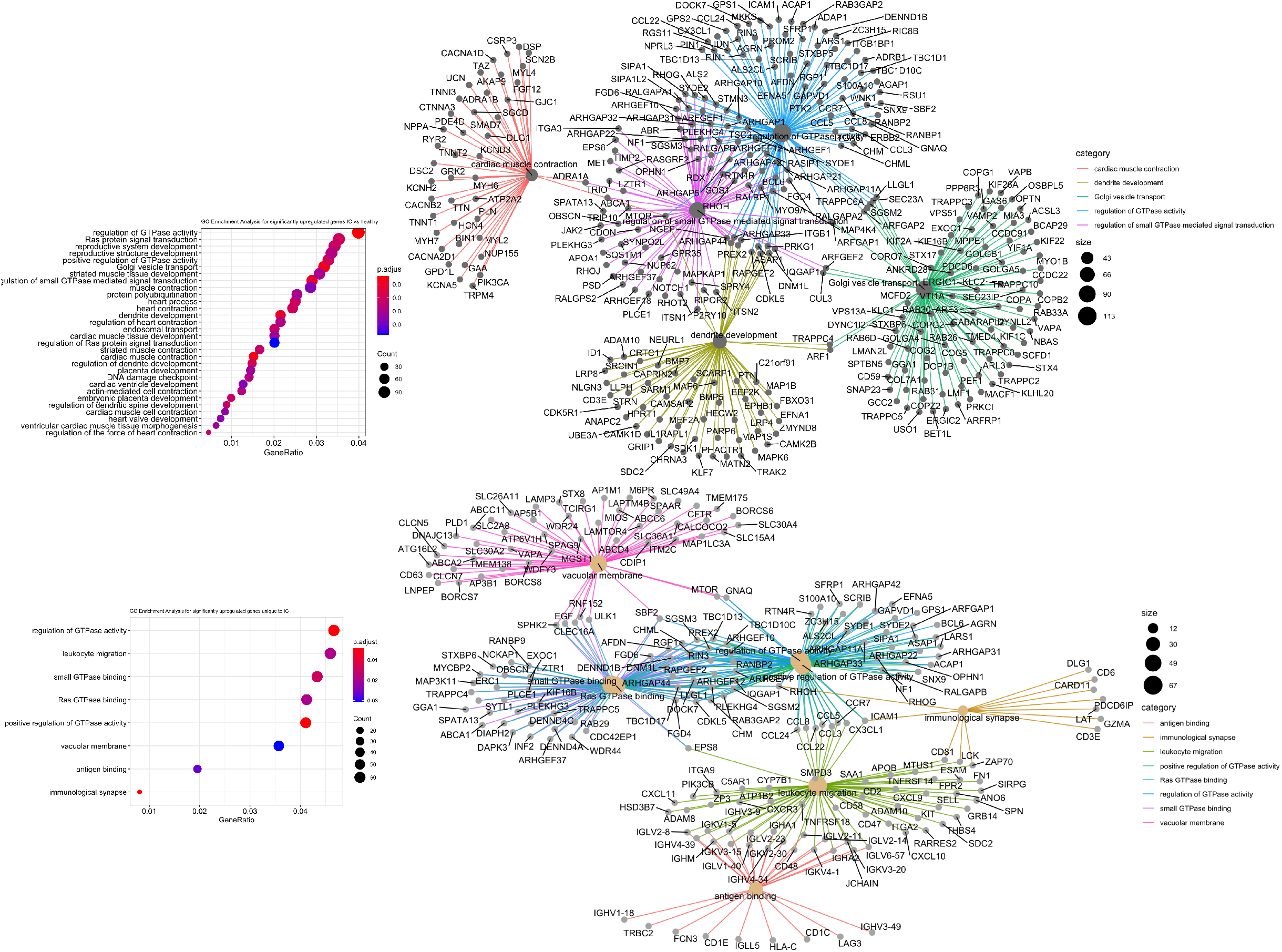
Enrichment analysis for all significantly changed genes in the IC (ischemic cardiomyopathy) vs Healthy contrast group where graph A shows enriched cellular processes and map B provides visualisation of the top highest ranking processes and corresponding genes. Gene set size that was enriched and enrichment p-adjusted value provided with the plots. Panels B and C represent gene enrichment for unique and significantly changed genes in the IC vs Healthy group.

### Human left ventricular tissue proteome analysis highlighted underlying metabolic differences in ischemic and hypertrophic heart states

Correlation between the expression levels of mRNA and protein is relatively difficult to establish with poor predictive power for the protein levels based on the gene expression^65–67^. Such correlation variability has been attributed to different regulatory mechanisms affecting transcriptional and translational processes^87–89^. Despite that, it was necessary to establish if proteome from a myocardial tissue rich left human ventricle could complement RNA-seq data.

While investigating protein abundances (Fig. 7), it became clear that the samples were quite similar as was the case with RNA-seq data (Fig. 2). Protein abundance values revealed a relatively homologous protein expression pattern across all samples as evident from the clustering based on the correlation values between sample protein levels (Fig. 7, A). Despite the high degree of shared features, sample similarity evaluation (Fig. 7, B) allowed the emergence of sample groups for dilated and ischemic cardiomyopathy as well as non-failing heart.

**Figure 7.**
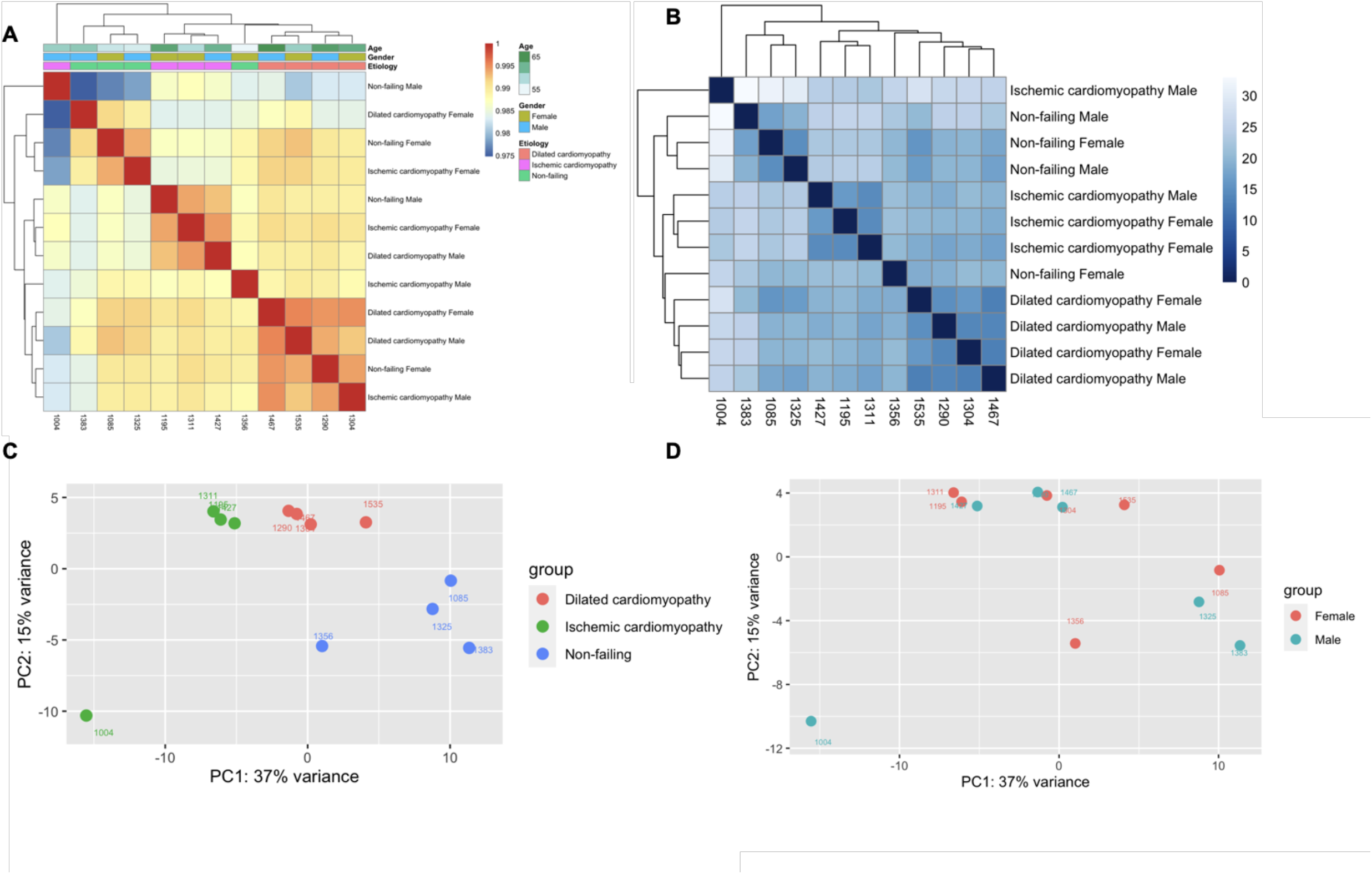
Human left ventricle bulk proteome (PXD008934) abunace clustering and distribution analysis showing Spearman correlation calculated distances (A) and euclidean distances (B) for rlog transformed abundance values (LFQ) using complete-linkage hierarchical clustering method; sample distributions across top two principal components are shown in the PCA plot grouping by condition (C) and gender (D).

Further exploration indicated that different pathological state samples vary less with clear separation between ischemic and hypertrophic conditions as well as healthy tissue. In addition, there were no gender dependent effects for the observed clusters (Fig. 7, C&D).

The next step of the analysis was to investigate for protein enrichment and compare the corresponding gene values with the data from RNA-seq study. Proteome data had a substantially lower recovery of data points (close to 3,000) when compared to nearly 19,000 for RNA-seq (Fig. 3; Supplementary Fig. 9). As expected, heart dilation leads to not only increased strain over heart but also causes subsequent muscle tissue remodelling (Fig. 9, A). There were 13 genes that showed a significant change in the RNA-seq samples as well as their matching counterparts on the protein level in the same contrast category (Supplementary Table 5&6). For example, Natriuretic peptides precursor A (NPPA), Aortic Carboxypeptidase-Like Protein (AEBP1) and Collagen Type XIV Alpha 1 Chain (COL14A1) genes as well as their corresponding proteins showed a significant upregulation in dilated cardiomyopathy; in a similar fashion Myosin heavy chain, α isoform (MYH6) and ADP-Ribosyltransferase 3 (ART3) were downregulated. All of these genes point to the remodelling events within the tissue, however, only several genes that showed enrichment on the protein level could be clustered based on their cellular role (Fig. 9, C).

Interestingly, Titin (TTN) expression levels dropped significantly but the reverse was true when evaluating for its protein levels (Supplementary Table 5&6; Supplementary Fig. 10). This bifurcation might likely occur due to multiple factors, namely mRNA stability and protein half-life^90^, which also demonstrates that gene or protein expression values cannot be used as sole measures but rather a systematic approach is needed.

A completely different picture can be seen when looking into ischemic heart transcriptome and proteome (Supplementary Table 7&8; Supplementary Fig. 10) and while functional enrichment in proteome study pointed toward lipid biogenesis and cellular respiration processes, the overlap between transcriptome and proteome only showed the enrichment for heart muscle hypertrophy, regulation of the heart rate as well as contraction force (Fig 8, B&D). Trying to compare protein vs gene expression further complicated picture (Supplementary Table 7&8; Supplementary Fig. 10) as there was less agreement in expression changes. As a case example, Myosin Heavy Chain 7 (MYH7) had a slight upregulation under ischemic conditions on the gene expression level but was markedly reduced on the protein level. This division in the expression values likely points to the complex regulatory mechanism for MYH7 under ischemic conditions as it is usually linked to the dilation and hypertrophy of the heart^91^. Several other genes, specifically Cytochrome C Oxidase Subunit 8A (COX8A) and Coenzyme Q-binding protein COQ10 homolog B (COQ10B), which are linked to the ischemic injury and loss of mitochondrial integrity^92,93^ showed similar patterns in gene and protein LFC.

**Figure 8.**
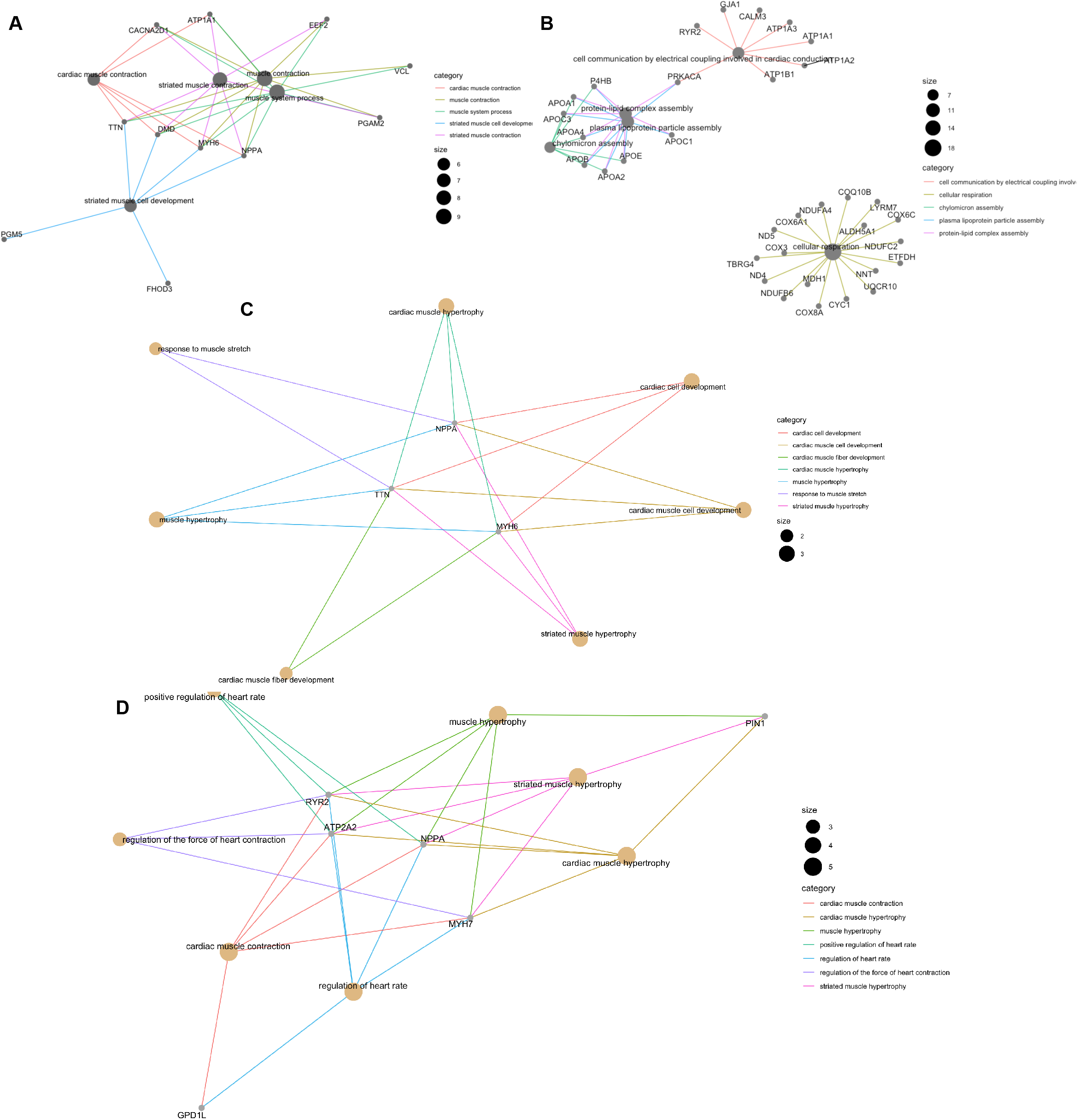
Enrichment analysis for all significantly changed proteins in DC (dilated) and IC (ischemic) cardiomyopathy when compared to healthy samples where A shows enriched cellular processes for DC vs healthy and B provides visualisation for IC vs healthy in human left ventricle proteome. Panels C and D show gene names that are shared between significantly changed proteome and transcriptome (PRJNA477855) for DC and IC, respectively.

Reverse was true for some of the genes that are reported to be involved in heart failure Kininogen 1 (KNG1)^94,95^, Retinol Binding Protein 4 (RBP4)^96^, Apolipoprotein B (APOB)^97^(Supplementary Table 7&8; Supplementary Fig. 10). This time no immune system associated enrichment was found for myocardial tissue rich samples as compared to RNA-seq data (Fig. 6) which likely confirms the complex composition of heart cellulome and presence of other cells that might be infiltrating tissues at different time points as a disease progresses. Overall, enrichment data (Fig. 8, B&D) for iscemic cardiomypathy demonstrated metabolic changes involving lipid generation and other proteins responsible for cellular respiration integrity.

### Single cell RNA-seq analysis of mice heart tissues reveal intricate cellulome composition that shared definitive markers with human heart RNA-seq data

To better appreciate the cellular composition of the heart, a single cell study on the murine non-myocyte cardiac cellulom was analysed and integrated with earlier studies.

While differences between species is a hurdle, this initial analysis aimed to get a better understanding of what cells can be found in the heart, their relative proportions and marker genes, and compare all of that with the findings in the human heart samples. Mouse heart preparations with cardyomyocyte cell population mostly removed (Fig. 9) split the remaining cells between matrix fibroblasts and subtypes of fibroblasts (the largest proportion), as well as various types of lymphocytes and leukocytes. Several interesting subgroups, for example Axin2+ cells, displaying stem-like cell properties and involved in fibrotic and regenerative events^98^ were found. Comparative analysis between human bulk RNA-seq significantly changed genes (either up-or down-regulated) and mouse single cell RNA-seq markers revealed a number of matches (Table 1). Most notably, dilated cardiomyopathy conditions were predominated by cardyomyocites and fibroblast like cells with some immune cell types. This was reversed in ichemic conditions with a high immune cell infiltration (Table 1). Comparing how different and non-cardiomyocte enriched cells cluster in mice heart tissues (Fig. 10&11), we can see that the separation was quite distinct where marker gene patterns (Supplementary Fig. 11) allowed to differentiate this rich cellulome. Cross-referencing single cell sequencing data with proteome analysis as well as bulk RNA-seq data (dilated cardiomyopathy) revealed two genes, ART3 and microfibril-associated glycoprotein 4 (MFAP4) to be also matched with mice heart cellulome markers. ART3 has been reported previously to be expressed in the heart^99^ but MFAP4 has several strong links to heart hypertrophy ^100^. Ischemic heart dataset analyses did not reveal such an overlap between bulk- and single-RNA-seq datasets as well as proteome analysis.

**Figure 9.**
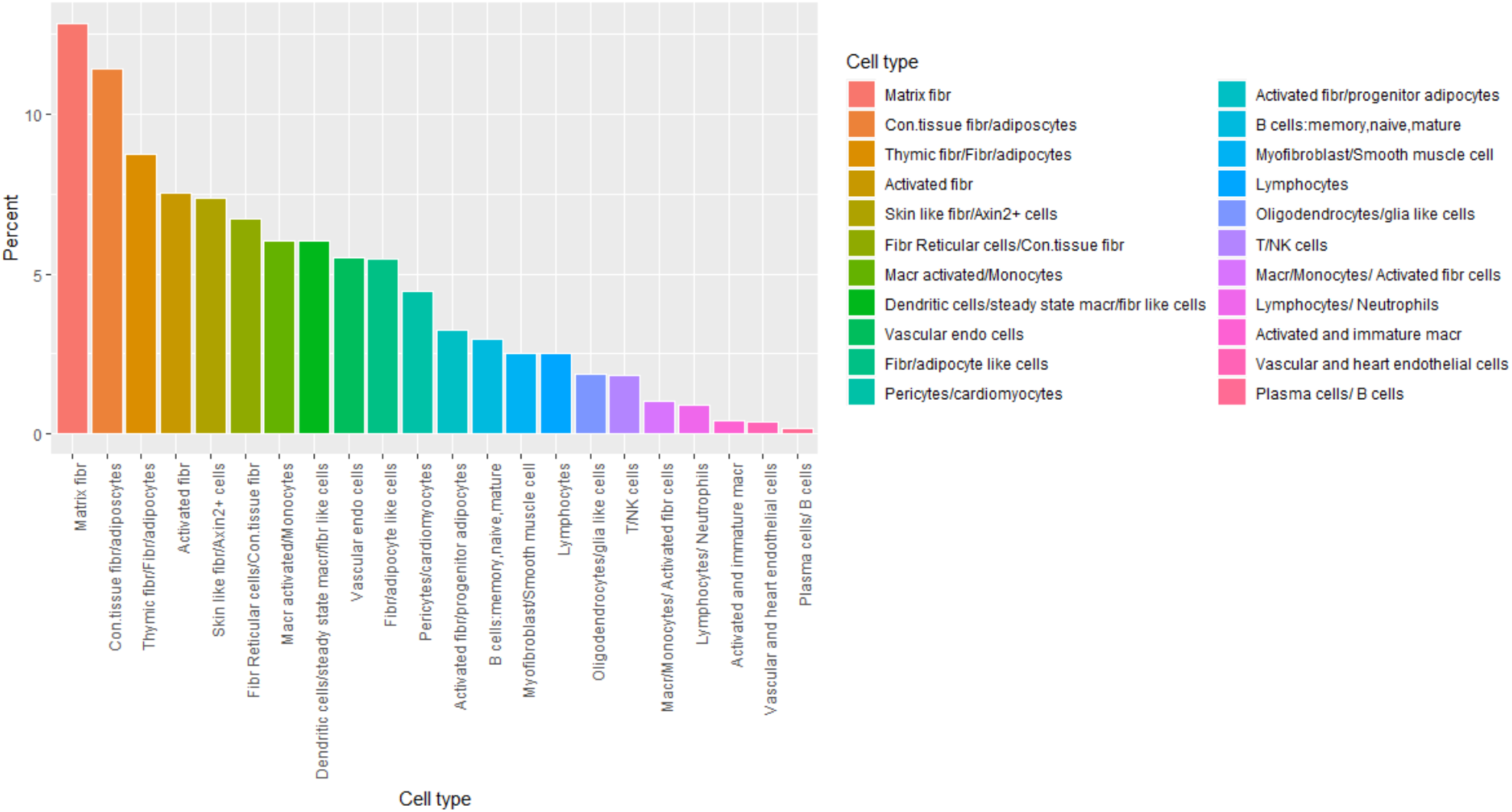
Mouse non-cardiomyocyte single cell RNA-seq (E-MTAB-6173) cellulome composition. *Some longer names were abbreviated; for full names, please refer to Supplementary Table 9.

**Figure 10.**
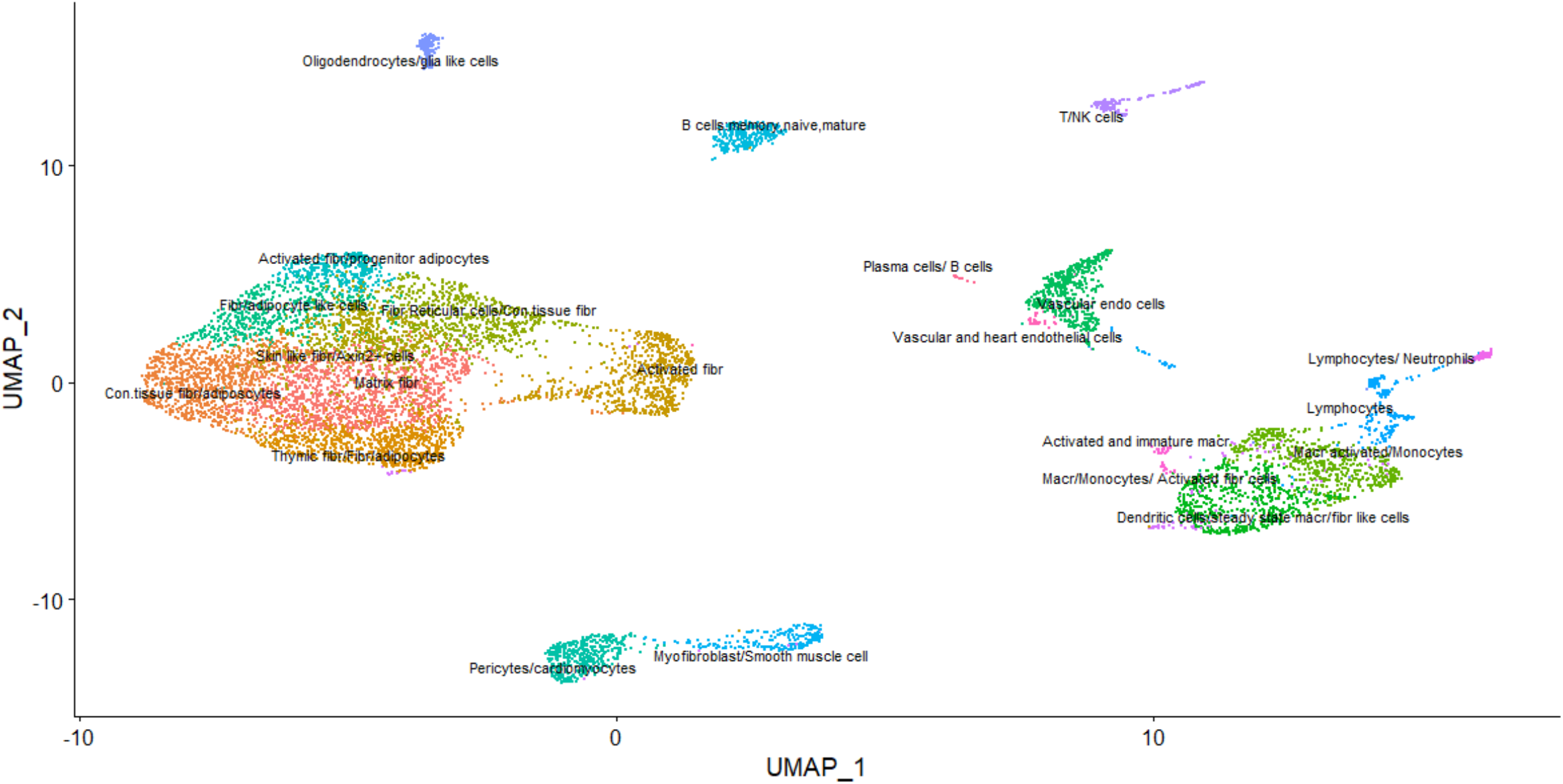
Mouse non-cardiomyocyte single cell RNA-seq (E-MTAB-6173) cellulome UMAP decomposition showing relative distances and the uncovered clusters of different cells. *Some longer names were abbreviated; for full names, please refer to Supplementary Table 9.

**Figure 11.**
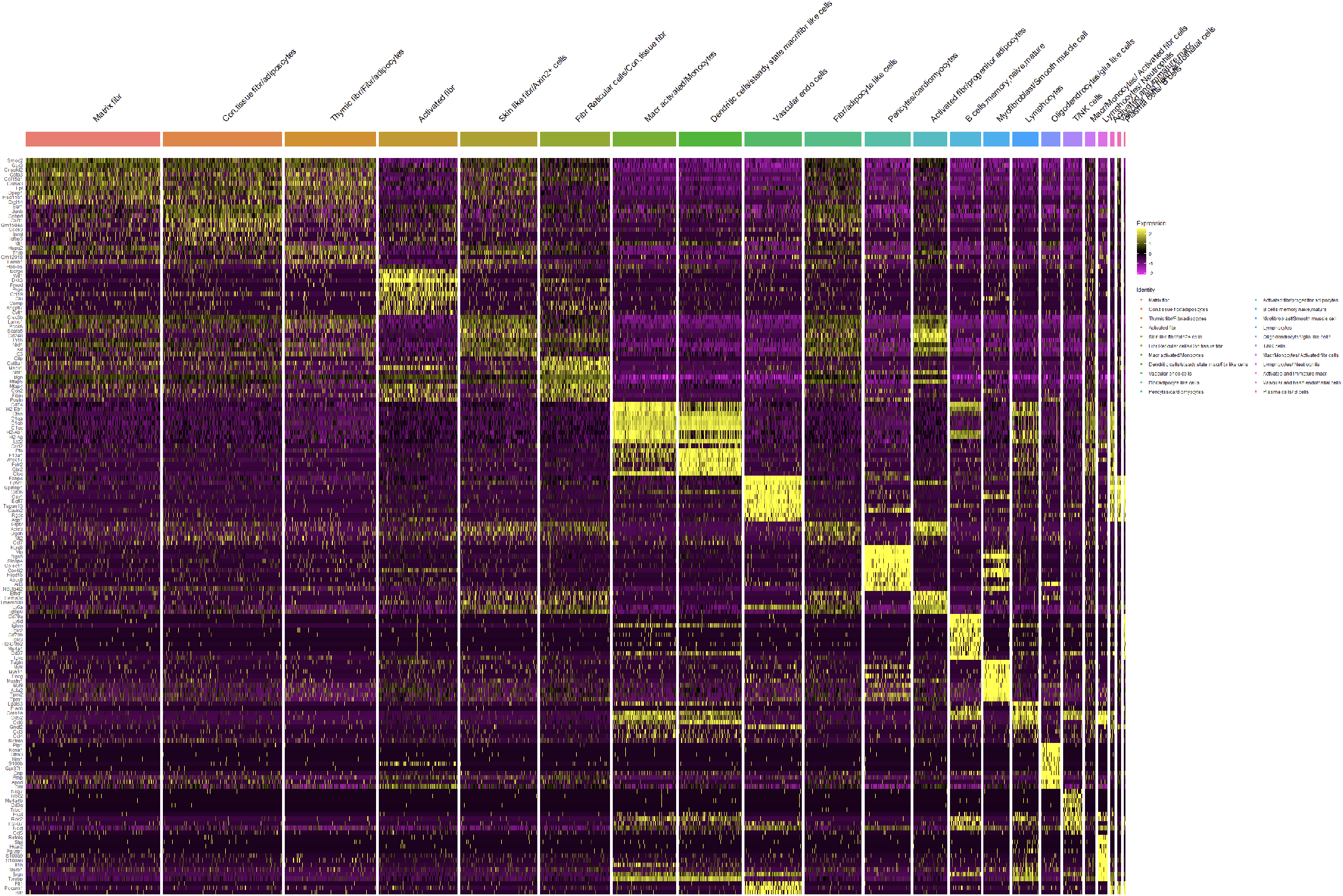
Mouse non-cardiomyocyte single cell RNA-seq (E-MTAB-6173) cellulome marker gene heatmap for the uncovered clusters of different cells. *Some longer names were abbreviated, for full names; please refer to Supplementary Table 9.

### Single cell RNA-seq analysis of the human heart left ventricle indicated the existence of divergent cell types for hypertrophic and ischemic tissue conditions

Single cell sequencing of human left ventricle revealed a complex mixture of cell types with expected cardiomyocytes, myoblasts and heart smooth muscle cells comprising nearly 65% of all cells and lymphoid cells adding up to more than a quarter of all cell populations combined (Fig. 12&13). These observations confirmed earlier findings (Fig. 5&6) where gene expression patterns suggested the involvement of both immune and other cell types that might contribute to fibrotic and remodelling events within the heart tissue. Specific marker genes for the human left ventricle showed varying expression patterns but a clear distinction between cardiomyocytes, heart smooth muscle cells or myofibroblasts required an elaborate combination of multiple marker genes to differentiate the groups precisely (Supplementary Fig. 12&13).

**Figure 12.**
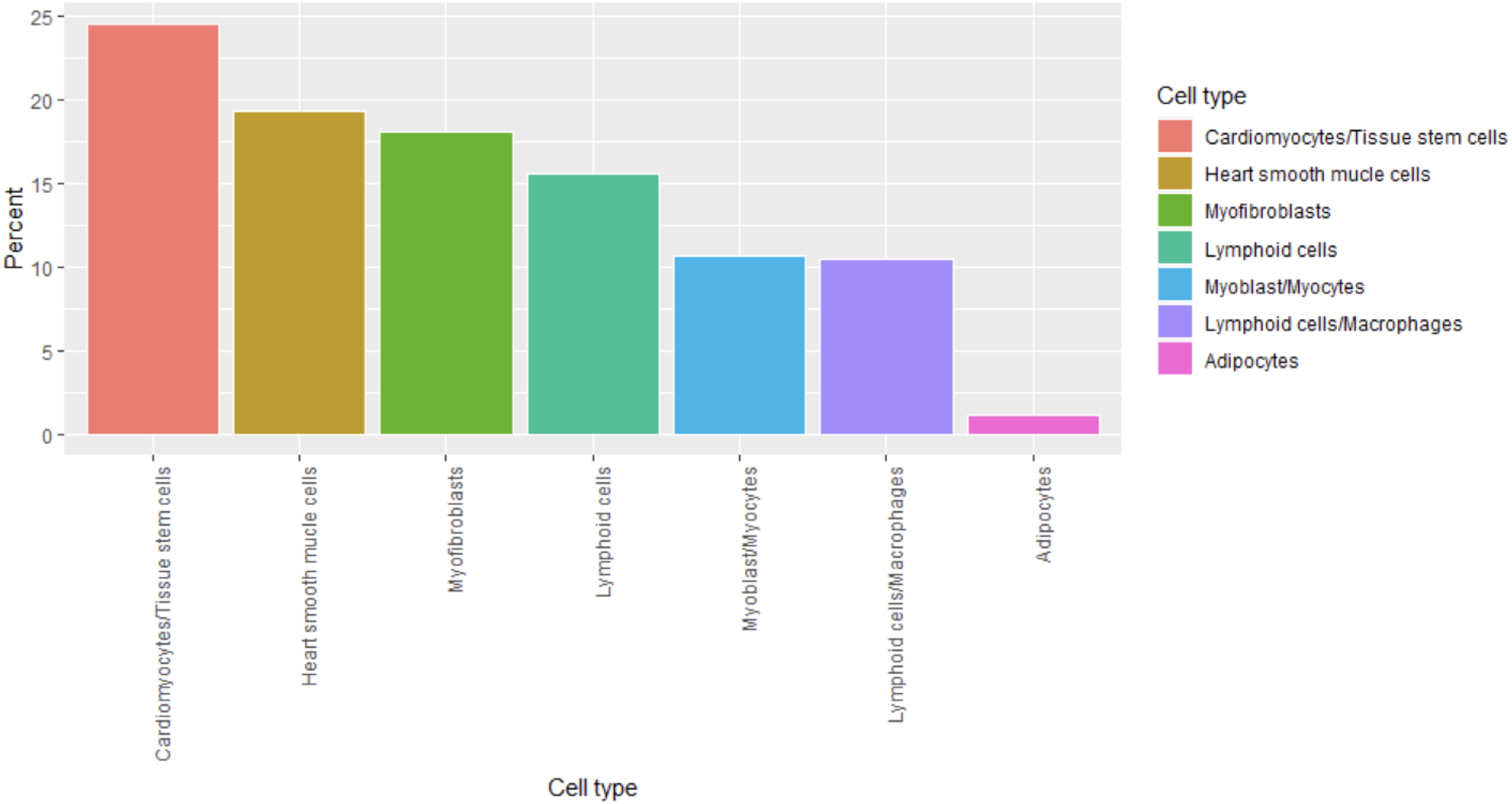
Human heart left ventricle cellulome composition.

**Figure 13.**
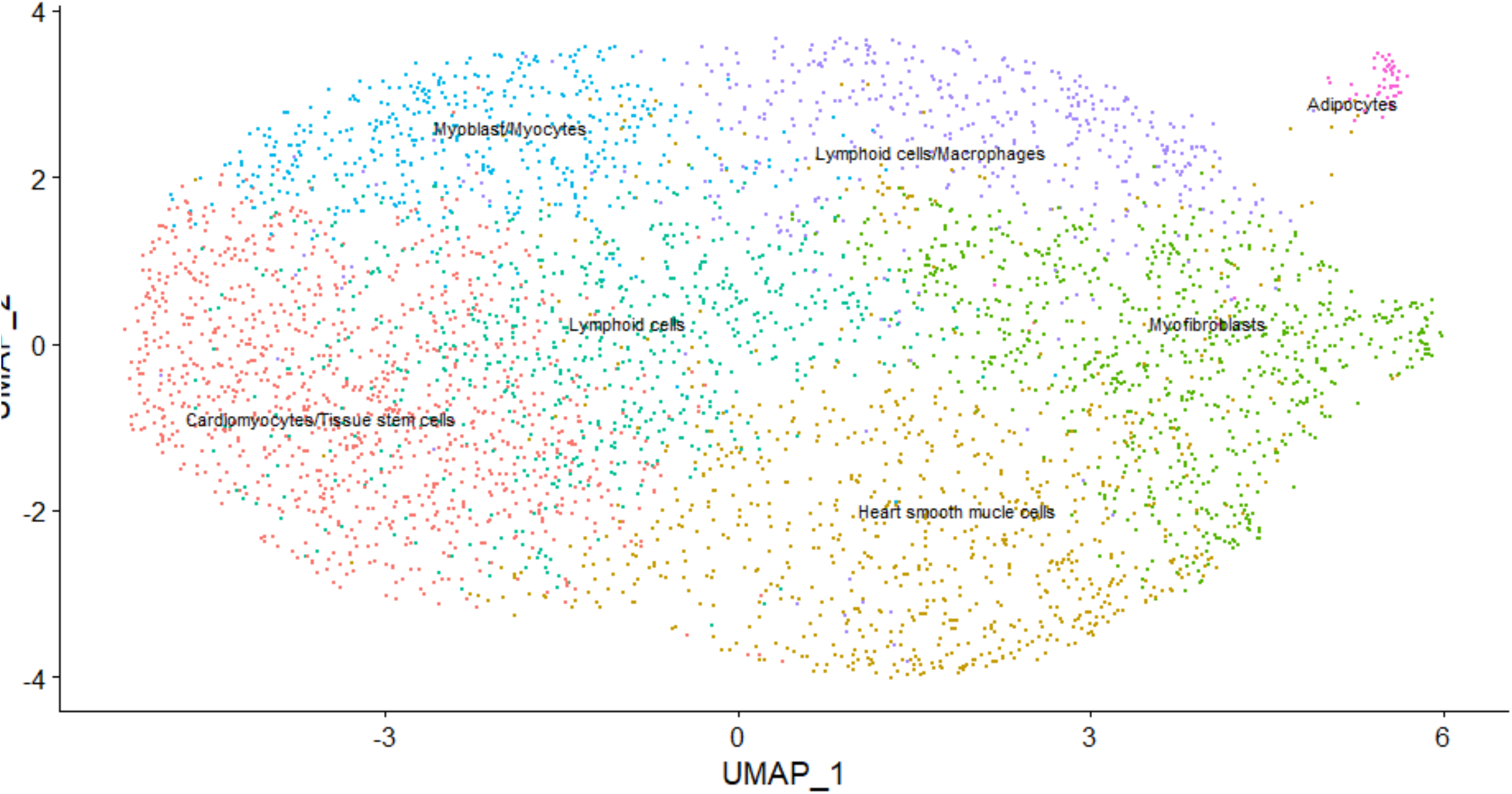
Human heart left ventricle cellulome UMAP decomposition showing relative distances and the uncovered clusters of different cells.

Further exploration of the dilated cardiomyopathy genes that changed significantly and had corresponding markers in human left ventricle bulk RNA-seq identified a change in the expression for genes likely involved in heart tissue remodelling; however, when this set was cross-referenced with matching proteome analysis, it did not return any hits.

Haemoglobin subunit alpha and beta (HBA1/2,HBB) was significantly upregulated in human bulk RNA-seq (contrast: DC vs Healthy) but showed a moderate change in single cell myocyte/myoblast population when this cell group was compared against the rest of heart cells (Table 1). Alpha subunit expression of the said globins has been implicated in the vascular tone and function maintenance^101,102^; together haemoglobin expression might suggest a compensatory mechanisms for the tissue undergoing contracticle and remodelling stress. Similarly, Orphan nuclear receptor 4A1 (NR4A1) showed a marked upregulation in the hypertrophic state while in a healthy left ventricle it was low (Table 1). NR4A1 has recently been described to play a role in cardiac stress responses and hypertrophic growth^103^. As a complete contrast, dilated cardiomyopathy showed a marked loss in adipocyte signatures (Table1), for example, fatty acid-binding protein 4 (FABP4) has been shown to contribute to cardiac metabolism^104,105^; thus, observed alterations might indicate a change in the energy metabolism of the heart.

Interestingly, ischemic heart tissue was very similar to the hypertrophic state when compared to human left ventricle cellulome. For example, a notable upregulation in most of the genes HBB, HBA1/2 and NR4A1 was matched between the different pathological states; however, Lumican (LUM) and HBB were also found to be significantly upregulated in ischemic heart proteome analysis. LUM has been shown to propagate the pro-fibrotic events in the heart failure^106^. The downregulated genes in ischemic conditions also followed similar patterns to the hypertrophic heart observed earlier, notably, FABP4 and Glycerol-3-Phosphate Dehydrogenase 1 (GPD1) belong to the gene group involved in lipid and amino acid metabolism^107,108^. While ischemic cardiomyopathy downregulated genes are dominated by adipocyte associated markers in this cross-reference analysis, TTN was also found to be downregulated under Myoblast/Myocytes group. TTN has been linked to remodelling and changes in the ischemic heart^109^ which based on the present study findings could be used to differentiate between hypertrophic and ischemic changes.

### New scoring system to evaluate genes using a two-step machine clustering approach revealed sets of disease specific interactors

The main challenge of biological data integration is to assign appropriate weights when combining multiple data sources in order to avoid biases. As previously shown, the richest data available is from bulk RNA-seq experiments (nearly 19,000 data points) while single cell RNA-seq and proteome analysis offers much less coverage and depth (close to 3000 data points). Thus, a decision was made to devise a scoring system that would take the advantage of bulk RNA-seq data and match with the data mined from multiple resources. Single cell or proteomics data had a very small overlap with bulk RNA-seq data (Supplementary tables 5–8); moreover, regular RNA-seq is still a preferred choice for multiple research studies and a more universal scoring system for this type of data is needed when the goal is to find specific genes that may play a role in a disease based on their expression characteristics. As a result, this study also aimed to introduce a score that could be supplied to machine learning pipelines to group existing data points and potentially predict genes with similar expression as well as disease causality patterns. The core of the equation (eq. 1) is the scaling of LFC value for a given contrast (e.g., disease vs healthy state) by a total association score retrieved from the Open Targets platform^91,110–113^. The score takes into consideration multiple data resources and evidence for a given gene, that is, whether there is a clinical precedence, reports in literature and/or known interactors as well as therapeutic compounds, all of this is combined into a single value which is between 0 and 1 where the higher value, the more probable association between the gene and a specific disease. 229 associations were retrieved for ischemic cardiomyopathy and a far larger number of gene scores (3521) was downloaded for dilated cardiomyopathy^44,114^. Moreover, using LFC as a base score allows to forgo the need to account for the different sequencing platforms or sequencing depths as it is a dimensionless measure showing the direction of the change for a specific expression.

To try and identify potential links between significantly changed genes in a given contrast a two-step machine learning approach was employed using Gaussian mixture models to identify gene clusters with the highest probability to share similar expression patterns and subsequently, each cluster can be further analysed using agglomerative hierarchical clustering to achieve a better refinement between associations. Before proceeding with the analysis, additional resource, STRING database, was parsed to retrieve the number of protein-protein interactions for a set of genes of interest. Each gene’s translational product – protein, shares a network with other proteins where the size of such a network might be varying. To estimate the impact of expression changes as evaluated by LFC Score, an assumption was made that if a protein is known to have multiple interactors, then it is likely that more pathways and/or processes will be perturbed compared to a smaller and isolated network. It is is important to note, that the applied machine learning approach takes into account all the features that a gene has and by performing advanced clustering it is possible to identify expression groups with new and interesting patterns that can be probed further. For example, a gene cluster that had a substantial change in expression, has known links to a disease and belongs to a larger network might be of interest when selecting potential targets. This, however, does not mean that the identified group will always belong to the same network; the identified grouping specifically differentiates genes and allows to select targets which may have multiple other interactions, change significantly and have stronger or weaker associations to a disease.

Gaussian mixture model based clustering revealed approximately the same number of features across dilated and ischemic cardiomyopathy groups (Fig. 14). To test, the impact of LFCChange Score the analysis was compared with a regular LFC. In the case of dilated cardiomyopathy there was a notable difference in the identified cluster distributions; in contrast, ischemic cardiomyopathy did not show such a noticeable difference primarily because the association scores were few and very low for this cardiopathology (Ischemic cardiomyopathy mean for association score: 0.00023; Max value: 0.01960; dilated cardiomyopathy mean for association score: 0.07050; Max value: 1). It became apparent that the more associations can be used as weights, the better resolution in clustering can be achieved.

**Figure 14.**
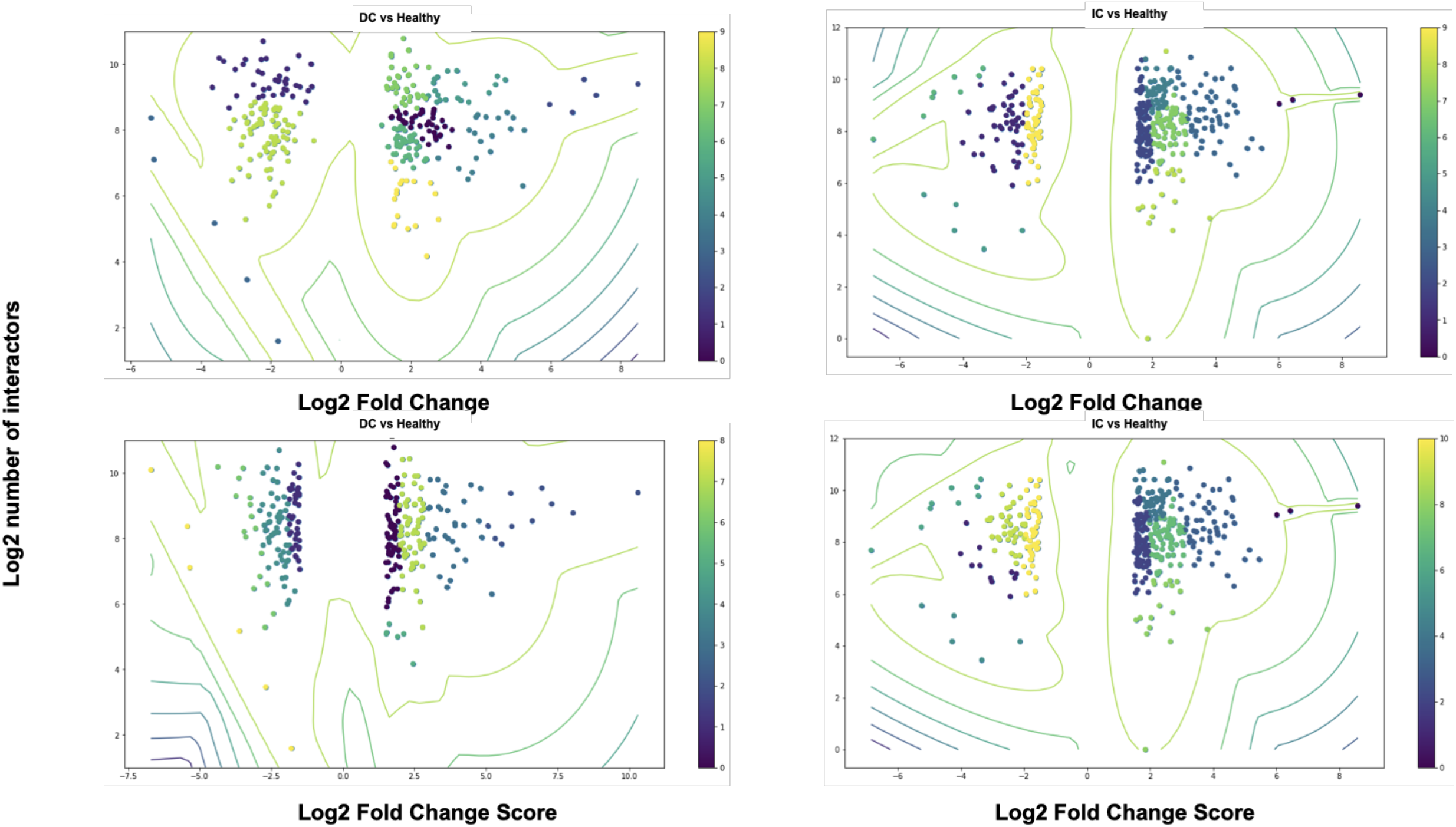
Human heart left ventricle bulk RNA-seq GMM clustering showing specific grouping based on either Log2 Fold Change or Log2 Fold Change Score against known or predicted number of interactions for that gene. Clusters investigated in the following conditions DC (dilated cardiomyopathy) vs Healthy and IC (iscemic cardiomyopathy) vs Healthy. Colour bar shows the specific cluster number and colour association.

This exploration was followed by the extraction of identified clusters and a downstream hierarchical clustering to demonstrate that identified groups can be sub-divided into finer sections based on each gene’s parameters (Fig. 15). For example, gene set from one of the bigger GMM clusters - cluster 0, (Fig. 14) for dilated cardiomyopathy was probed further to reveal subtle variations between genes. A case example of one of the sub-clusters, Mothers against decapentaplegic homolog 7 (SMAD7), Syntaxin-1B (STX1B) and Transcription factor SOX-17 (SOX17), show how genes with similar expression and network profile can be grouped and in this case, these genes belong to different branches of a complex network regulating tissue morphogenesis, vesicle docking and growth. Some of the ischemic cardiomyopathy sub-clusters, such as Tumor necrosis factor receptor superfamily member 11B (TNFRSF11B) and Serine/threonine-protein kinase pim-2 (PIM2) showed convergence of two signalling branches via Myc proto-oncogene protein (MYC). While capturing different gene sub-clusters that show similar characteristics based on the classifiers used does not guarantee that they will belong to the same pathway or network, this approach could still aid in building new networks for downstream analyses and experimental validation when identified genes from a cluster are used as a seed points.

**Figure 15.**
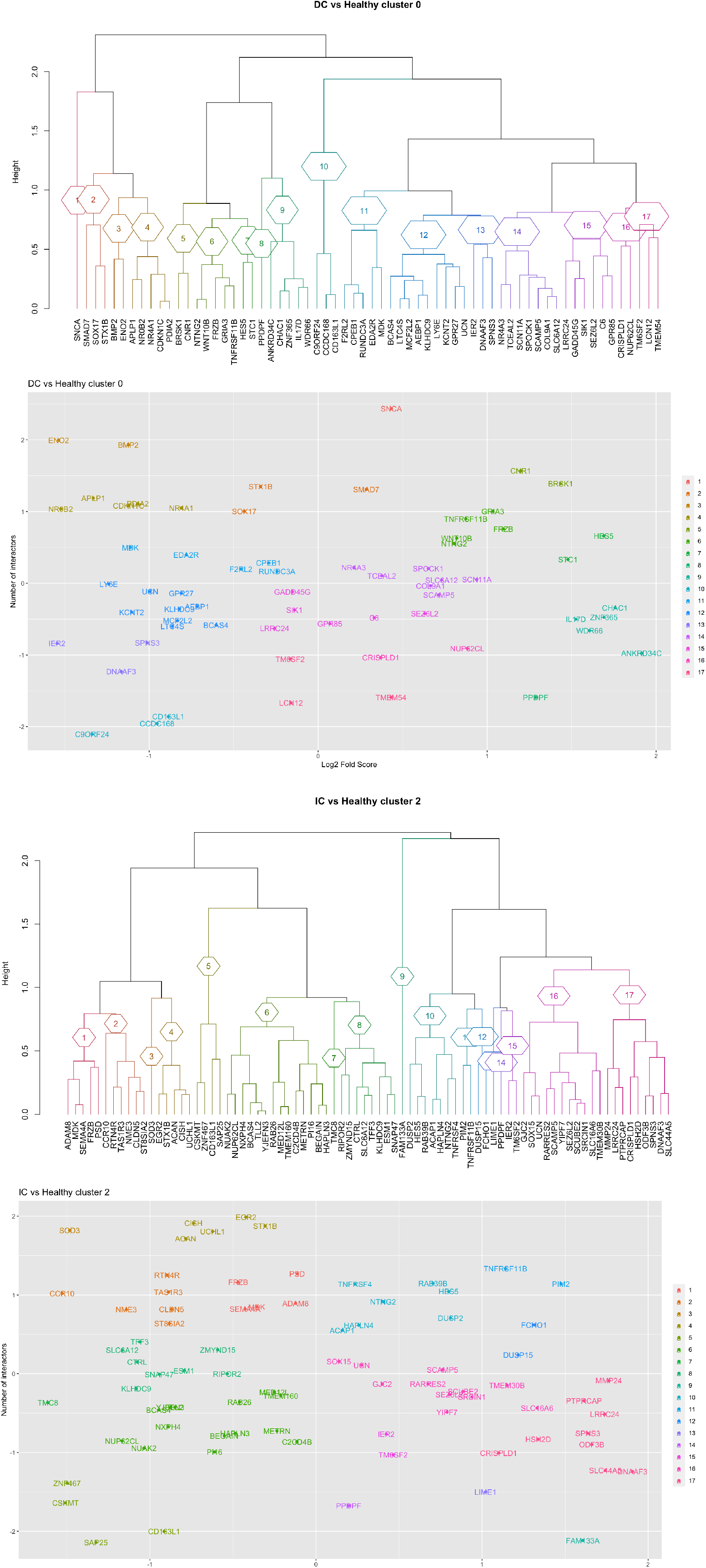
Human heart left ventricle bulk RNA-seq GMM analysis identified multiple clusters which were further subjected to hierarchical clustering (dendrogram panels). Representative GMM cluster hierarchical sub-clustering is shown with each dendrogram leaf labeled based on a specific sub-cluster number. Respective right panels show grouped gene distribution for z-score scaled parameters on which the sub-clustering was performed DC (dilated cardiomyopathy) vs Healthy and IC (ischemic cardiomyopathy) vs Healthy contrasts are shown.

Another exploratory possibility was to extract genes from GMM clustering that are shared between dilated and ischemic cardiomyopathy (160) and perform hierarchical clustering on the set by juxtaposing two clusters (Supplementary Fig. 14). Some of the gene groups (coloured branches) are matched between two pathologies based on the gene disease association, LFC and the number of interactors. For example, such one shared group of Protein Wnt-9a (WNT9A), HBB and (F-box and leucine rich repeat protein 16) FBXL16, belong to the same large network of interactors likely playing a role in tissue function and local signalling events. The described analysis could be very useful in understanding potential convergence points for diseases and how shared genes are grouped per disease profile.

Finally, machine learning approaches rely on the breadth and quality of data. This is especially evident in the case of hypertropic and ischemic condition derived gene clustering (Fig. 14) where the richer set of information can help in differentiating between clusters better; thus, the reported methodology could be used with the in-house data to enrich for known associations and provide more information for scoring algorithms.

## Discussion

Dilated cardiomyopathy is an important cause of heart failure characterised by the ventrical enlargement and subsequent systolic dysfunction. In contrast, ischemic cardiomyopathy is a clinical manifestation with a complex causality ranging from coronary artery disease to other changes in the heart muscle that decrease nutrient and oxygen supply. A wide spectrum of etiologies, including inherited, inflammatory and/or infectious diseases can predispose the heart to this pathological remodelling^44^.

Studying the cardiac impairment resulting from heart dilation or ischemia is complicated by the mixture of known as well as idiopathic causes. Moreover, integrating a complex transcriptional landscape might be difficult as evident in the past studies reporting on experimental or meta-analyses^4,11,115,116^, this is because collected tissues for experiments differ to some extent and a sample population might introduce various other cofounding factors (e.g., past or current treatment, body metabolic state, etc.). In addition, depending on the statistical assumptions made and model selection results may vary. Thus, one of the aims of this study was to explore the different *omics* resources for cardiomyopathy and what we could learn from integrating such datasets.

The first part of the analysis focused on the human left ventricle tissue bulk RNA-seq analysis for two indications: dilated and ischemic cardiomyopathy. By analysing significantly changed genes, it was possible to see a subtle separation between hypertrophic and ischemic heart conditions. Fo example, dilated cardiomyopathy tissue had a number of significantly upregulated genes (BMP2, MYOZ1and ENO2) that showed strong associations with myocardial tissue remodelling and structural changes when compared to the healthy samples^55,56^. Some other genes, such as RPS17, SLITRK4 and GLT8D2, belong to newer additions of potentially valuable genes and just recently implicated pathways in dilated cardiomyopathy. These genes are involved in protein synthesis and post-translational modifications as well as cell growth control^59–61^. Opposing group of genes that were downregulated (CA11, ICAM3 and ELOVL2) hints at the metabolic perturbations^44,57,58^ spanning the spectrum from cellular respiration changes to potentially loss of the membrane integrity in the tissue that is actively being remodelled. When contrasting these findings with ischemic heart conditions, there was a notable change in the upregulation of the pro-inflammatory and pro-fibrotic genes. For example, CX3CL1 is an especially intriguing gene as it encodes an atypical chemokine which can exist in either a membrane-bound form or as a soluble chemokine; the membrane integrated form is largely expressed on endothelial cells in myocardial ischemia and heart failure^62–65^. Another candidate gene and potential biomarker of note, TMEM259, has some associations with ischemic conditions as well as ER protein degradation pathways^65^. A number of other genes: FOC, REC8, FHOD1 as well as TAOK1 and MINDY2, are involved in the modulation of cell proliferation, immune signalling and protein turnover^69,117^. These findings provided the first hints of potential exacerbation of the ER stress as well as inflammation induced fibrotic tissue damage propagating the ischemia cycle. In addition, gene expression changes for the broad spectrum of cellular metabolism and growth processes were more pronounced in dilated cardiomyopathy.

The differences between dilated and ischemic cardiomyopathy were further highlighted when clustering genes based on their ontologies and involvement in cellular processes and pathways. Myocardium remodelling, ventricular cardiac muscle tissue morphogenesis and muscle tissue development as well as other tissue structure and integrity related processes were enriched for dilated cardiomyopathy (Fig. 5, A&B). With a further refinement – uniquely and significantly changed genes showed a specific clustering under microtubule, myofibril, sarcromere and contractile fibre process group (Fig. 5, C&D). Some of those genes, MYL1, DNAH6, MYOZ1 and ACKR2 could be of a special interest as potential therapeutic targets or biomarkers because of their reported roles in heart muscle function. For example, MYL1, DNAH6 and MYOZ1 were named in various reports linking them to hypertrophy, changes in contractility and myocardium cell function^44,74–76,118^. Tissue overgrowth mediated by these genes could be targeted to reduce excessive strain on the myocardium in the early stages of the disease development. ACKR2 has been demonstrated to reduce inflammation and vascular remodelling after myocardium injury, this identified upregulation might indicate compensatory mechanism for the tissue remodelling^44^. Thus, enhancing or stimulating this protective signalling might be an interesting therapeutic option (Fig. 6, A&B; Supplementary Table 3).

In contrast to hypertrophic heart muscle, ischemic heart enriched gene networks had clear links to ER stress; for example, SMPD3, Transmembrane protein 175 (TMEM175), Epidermal Growth Factor (EGF) and APOB have been shown to lead to ER stress when their normal function is perturbed^74^. In addition, FPR2 as well as CX3CL1 interlink inflammatory processes with higher ER protein turnover burden^78,119–122^ (Fig. 6, C&D; Supplementary Table 3). Under ischemic conditions, perturbations in oxygen and nutrient supply as well as undergoing cellular stress can lead to mitochondrial and proteome stability changes which likely propagates fibrotic remodelling events^33,65–67,79^. Also another intriguing finding was a number of chemokine ligands (e.g., CXCL11, CXCL10 and CCL5), some chemokine receptors (e.g., CXCR3 and CCR7) as well as other markers, such as CD2, to be significantly changed under myocardial ischemia (Supplementary Table 4). This, however, might be likely attributable to the T-cells and other lymphoid cells tissue infiltration as can also be seen in a mouse left ventricle non-myocardial cellulome study. A significant proportion of fibroblast and fibroblast like cells can also be found in a healthy human heart (Fig. 12, Table 1) and this population under myocardial stress conditions can change its proportions further by propagating pro-inflammatory and pro-fibrotic environment. Moreover, normal subpopulations of immune cells identified in the heart, such as monocytes, macrophages, mast cells, eosinophils, neutrophils B cells and T cells, can also be activated and lead to a pro-inflammatory state^56,61,117,123^. These considerations need to be taken into account when analysing data at different resolution levels where different types of cells can show a varying degree of contribution in the bulk transcriptome.

This was especially evident when juxtaposing enriched protein groups and comparing them with the corresponding gene values from the data of RNA-seq studies. As proteome data had about six times lower recovery then bulk RNA-seq (Fig. 3; Supplementary Fig. 9), it became clear that it is only possible to identify genes and their networks that are above the detection level and show substantial abundance. Despite these limitations, important marker molecules associated with dilated cardiomyopathy were found; that is, NPPA, AEBP1, MFAP4 and COL14A1 were upregulated both on the gene and protein level. All of these genes and their readouts on a protein level could potentially be used as biomarkers since there is experimental and clinical evidence for their role in the dilated left ventricle remodelling^124–126^. MFAP4 was also matched to the mice heart cellulome fibroblast markers; this target is quite interesting as it reoccured in all three types of *omics* datasets and has not only several strong links to the heart hypertrophy but has also been investigated as a potential therapeutic target^127^.

In the case of downregulated genes and their proteins, MYH6 and ART3 form a unique group; while MYH6 mutations are linked to hereditary cardiomyopathies^101,102^, ART3 function remains to be defined but it was found in cardiac proteome profiling^128^. The present study also identified ART3 as a pericyte/cardiomyocite marker from a single cell study for mouse heart cellulome with most myocytes removed. This not only confirms that ART3 expression allows it to be associated with cardiomyoctes and differentiated from other cells but could also suggest that the reversal of this downregulation might be a new therapeutic opportunity. In addition, TTN had a contrasting pattern were gene expression levels were decreased and protein expression was upregulated (Fig. 9, C; Supplementary Fig. 10). TTN mutations are well-documented for dilated cardiomyopathy; however, while mutated and truncated TTN proteins lead to the disease parthenogenesis, the higher expression role is not clear^100^. In addition, TTN was also found to be of low expression under Myoblast/Myocytes group in the human left ventricle single cell RNA-seq (Table 1). It is possible to hypothesise that as heart muscle remodelling progresses some of the compensatory mechanisms might increase contractile fibre and associated protein production at the same time RNA expression levels drop by secondary regulatory mechanisms to reduce the protein production burden. More in-depth experimental studies investigating TTN and its expression dynamics are necessary to understand whether there is any prognostic or therapeutic value.

Ischemic heart transcriptome and proteome (Supplementary Table 7&8; Supplementary Fig. 10) overlap only showed the enrichment for heart muscle hypertrophy, regulation of heart rate as well as contraction force (Fig 8, B&D). For example, while MYH7, COX8A, COQ10B had a slightly increased gene expression, protein expression values were markedly suppressed. MYH7 is a well-known driver of cardiac tissue hypertrophy^91,110–113,129,130^; thus, lack of nutrients reaching heart might prevent tissue growth and dampen related pathways. Moreover, decrease in COX8A might be a protective mechanism to reduce oxidative metabolism^92,93^. However, other perturbations, such as the loss of COQ10B ensuring mitochondrial integrity^94^ likely overcome measures against oxidative stress leading to ischemic tissue injury propagation (Supplementary Table 7&8; Supplementary Fig. 10). Other gene products, namely APOB, RBP4 and KNG1, playing the role in the heart failure were overexpressed despite reduced mRNA levels which could give a glimpse into the perturbed energy metabolism and tissue blood perfusion^95^. This sharp contrast could hint toward potential therapeutic avenues to inhibit RBP4 based signalling and APOB induced ER stress that are likely contributing to further tissue injury and remodelling. While the heart left ventricle proteome did not capture strong immune associations as previously shown in bulk RNA-seq, it is noteworthy that lipid metabolism associated proteins had a clear presence (Supplementary Table 7&8; Supplementary Fig. 10). Moreover, cardiomyocytes are not a homogenous group of cells as can be seen in bulk and single cell RNA-seq, this is also true on the protein level where a small subset of proteins show variability between cardiomyocytes in a mosaic pattern and can likely be further altered under pathological conditions^96,97,123^.

In parallel, all of the above findings were also compared to the human left ventricle single cell RNA-seq. While the majority of cells were mostly cardiomyocytes and other muscle tissue cells, more than a quarter was comprised of various immune cells (Table 1, Fig. 12&13). One of the most interesting findings was a matched significant upregulation between bulk and single cell RNA-seq as well as proteome data that returned LUM and HBB genes for ischemic heart conditions. Experiments with LUM demonstrated its ability to increase the levels of lysyl oxidase, collagen type I alpha 2, and transforming growth factor-β1, and decrease the activity of the collagen-degrading enzyme matrix metalloproteinase-9, thus these pro-fibrotic events are associated with a higher potential for a heart failure^118^. Targeting LUM might help control the fibrotic tissue transformation in the heart and it could also be used as a prognostic marker. Yet, the expression dynamics of LUM are not entirely clear as more recent reports indicate that LUM might be involved in compensatory and counterbalancing functions during active heart failure^107^. Such contradictions, reaffirm the complexities of underlying pathology mechanisms and further research is needed to understand at what heart failure stages these expression changes occur and when it is best to have a pharmacological intervention. Furthermore, alpha subunit expression of the globins has been implicated in vascular tone and function maintenance^108^; thus, it is possible that beta subunit expression might be involved in similar compensatory mechanisms for the tissue undergoing ischemic stress but again further research would help to establish the therapeutic potential of HBB and other globins. Another interesting candidate target was identified for the dilated cardiomyopathy, namely NR4A1. This orphan receptor showed a marked upregulation in hypertrophic state while in a healthy left ventricle its expression remained low (Table 1). NR4A1 has recently emerged as one of the key players in cardiac stress responses and hypertrophic growth^103^. Also, hypertrophic tissue state showed a marked loss in adipocyte signatures (Table1) and some of the downregulated genes followed similar patterns in the ischemic heart observed earlier - notably, FABP4 and Glycerol-3-Phosphate Dehydrogenase 1 (GPD1). FABP4 has been suggested to influence cardiac size and myocardial function under pathological states^106^ and it might indicate changes in the energy metabolism of the heart^105^. GPD1 is known to play a role in oxidative stress responses as well as affect lipid and amino acid metabolism^103^, it is possible that the observed downward shift in GDP1 expression is a compensatory mechanism and could be a valuable marker.

All of these observations, clearly delineate the need to appreciate the different levels of *omics* datasets. While bulk and single cell transcriptome, proteome as well as epigenome analyses^55,56,61,64,100,131,132^ provide us with varying degrees of resolution in cases of complex tissue and more so, in cases of wide spectrum pathologies, it might become difficult to integrate such variable datasets. Thus, to address the main challenge of biological data integration a scoring system that would take the advantage of bulk RNA-seq data and match with the data mined from multiple resources for each gene was devised and introduced in this study. As demonstrated earlier, the richest biological data is still only available from bulk RNA-seq experiments (nearly 19,000 data points) and all other resources, such as proteomics or single cell RNA-seq, had only a very small overlap with the genes identified from bulk sequencing (Supplementary tables 5–8); moreover, regular RNA-seq is still a more universal research choice to untangle transcriptional profiles; as a result a scoring system was used to capture the level of gene expression change along with any mined associations for that gene so that it was possible to supply this information to machine learning pipelines and group existing data points to predict biologically meaningful genes expression patterns.

Two-step machine learning approach returned multiple subgroups which showed similar multi-profile characteristics, for example, SMAD7, STX1B and SOX17 not only belonged to the same sub-cluster (Fig. 15) but are also a part of a complex network regulating tissue morphogenesis, vesicle docking and growth in dilated cardiomyopathy significantly changed genes^106,133^. Similarly, TNFRSF11B, PIM2 converged via MYC in the network linked to angiogenesis and anti-apoptotic pro-growth effects in ischemic cardiomyopathy group (Fig. 15). While genes belonging to the same cluster hint toward interesting target candidates, they are not necessarily direct interactors. Moreover, as can be seen there is a certain degree of separation for these genes^45^ but it could be useful when either building network models and using these genes as a seed points or predicting the extent of local network perturbations when a gene in a cluster is targeted. Another application of this method is to cluster genes that are shared between two pathologies as was the case for this study and using that establish gene groups that showed similar profiles in different conditions. For example, WNT9A, HBB and FBXL16 were both clustered to the same group for dilated and ischemic cardiomyopathy when a pool of 160 shared genes was subjected to agglomerative hierarchical clustering. This could be very useful in understanding potential convergence points for diseases, establishing shared expression patterns and selecting therapeutic targets that are substantially unique.

## Conclusion

Current strategies to treat the heart failure mainly target symptoms based on the left ventricle dysfunction severity. There is a notable lack of systemic studies and available biological data for an in-depth analysis of heterogeneous disease mechanisms on the scale of genomic, transcriptional and expressed protein level. The growing need for a better disease management was the main impetus behind this study to investigate if the shift in the analytical paradigm can be implemented to focus more on network centric and data mining approaches. This study, for the first time, demonstrated how bulk and single cell RNA-sequencing as well as proteomics analysis of the human heart tissue can be integrated to uncover heart failure specific networks and potential therapeutic targets or biomarkers. Most importantly, it was shown that transcriptomics data can be enriched with the minded data from public databases and used to identify specific gene expression profiles. To achieve this specific goal, a two-step machine learning pipeline was introduced. The described methodology could be very beneficial for the target selection and evaluation during the early stages of therapeutics development. Finally, the present study shed new light into the complex etiology of the heart failure differentiating between subtle changes in dilated and ischemic cardiomyopathy on the single cell, proteome and whole transcriptome level.

## ABBREVIATIONS

CVD: cardiovascular disease
DC: dilated cardiomyopathy
GMM: Gaussian mixed models
IC: ischemic cardiomyopathy
RNA-seq: RNA sequencing

