## Supplementary material for "Novel insights into potential therapeutic targets and biomarkers using integrated multi-*omics* approaches for dilated and ischemic cardiomyopathies": suplementary_materials

**Authors contributions**
AK devised the methodology, performed the analyses and wrote the manuscript; NB provided critical review, medical perspective and suggestions for the manuscript

**CORRESPONDENCE:**
Austė Kanapeckaitė,

**KEYWORDS:**
Target identification; Biomarker discovery; Dilated cardiomyopathy; Ischemic cardiomyopathy; Omics data integration; Machine learning for target prediction

**ABBREVIATIONS:**
CVD – cardiovascular disease; DC – dilated cardiomyopathy; GMM – Gaussian mixed models; IC – ischemic cardiomyopathy; RNA-seq – RNA sequencing

**HIGHLIGHTS:**
- First report of an integrated multi-omics analysis for dilated and ischemic cardiomyopathies.
- Identification of metabolic and regulatory network differences for the two types of cardiomyopathies.
- Introduction of a new scoring system to evaluate genes based on the size of their network and disease association.
- Two-step machine learning pipeline to uncover potential therapeutic target clusters.

**CONFLICT OF INTEREST:**
Authors declare no competing interests.

### Supplementary Figures

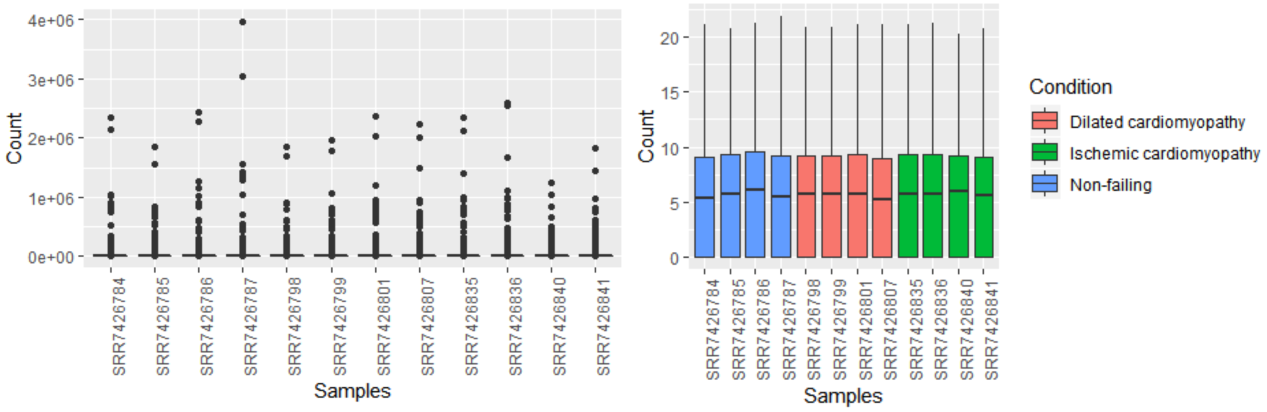


**Supplementary Figure 1.** Raw (A) and log2+1 normalised (B) sample count distributions for human left ventricle bulk RNA-seq (PRJNA477855).

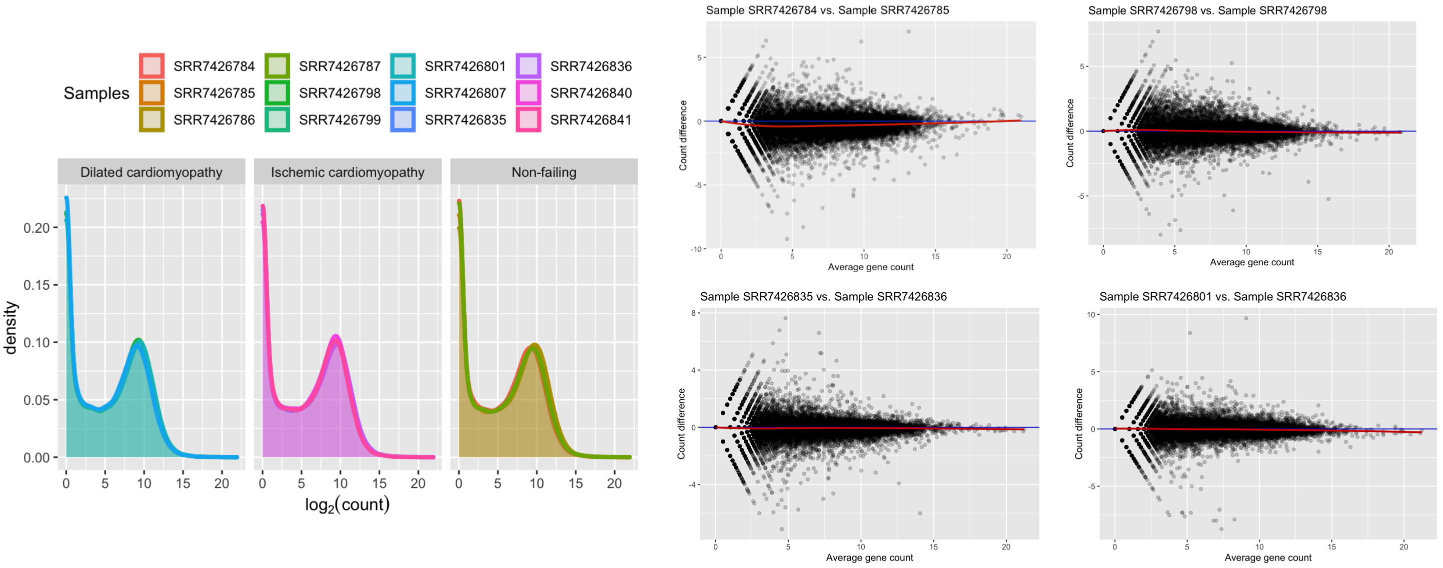


**Supplementary Figure 2.** Density distribution of log2+1 transformed sample counts (A) and MA plots for log2+1 transformed counts (B) where the red line indicates average differences.

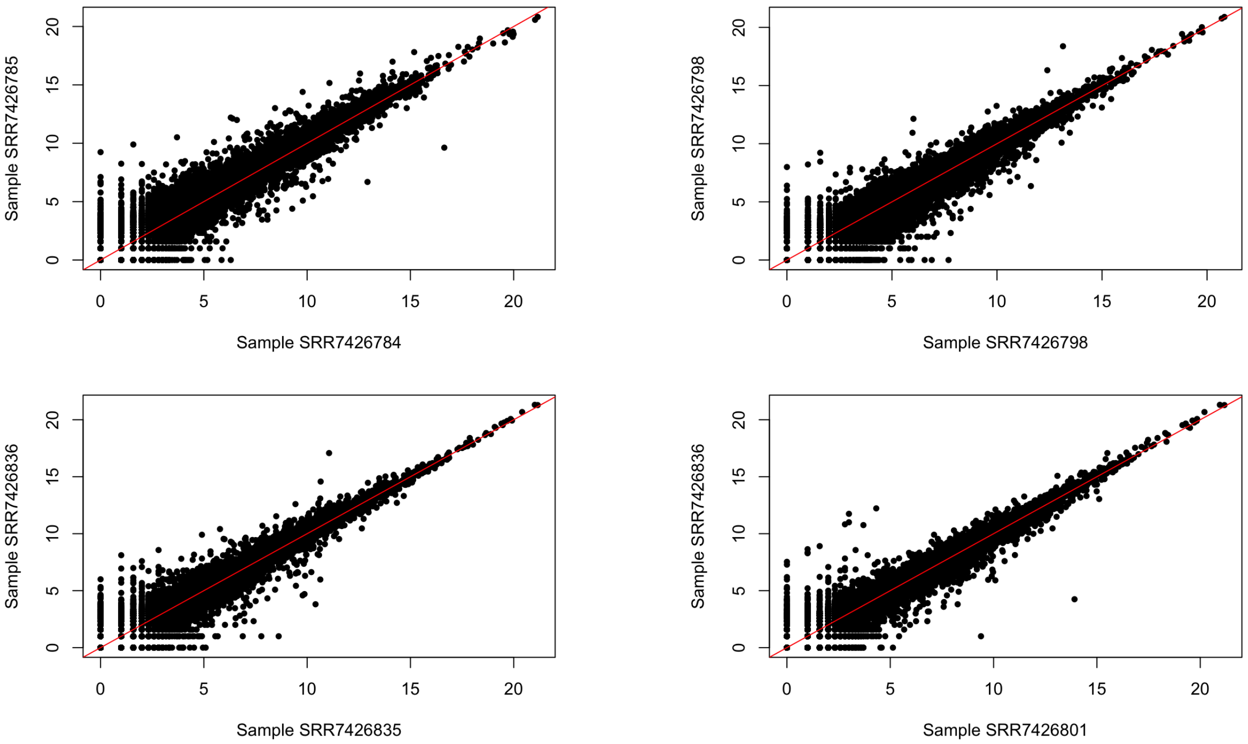


**Supplementary Figure 3.** Log2+1 transformed counts for samples plotted against each other to evaluate count distribution and sequencing depth. Representative combinations are shown.

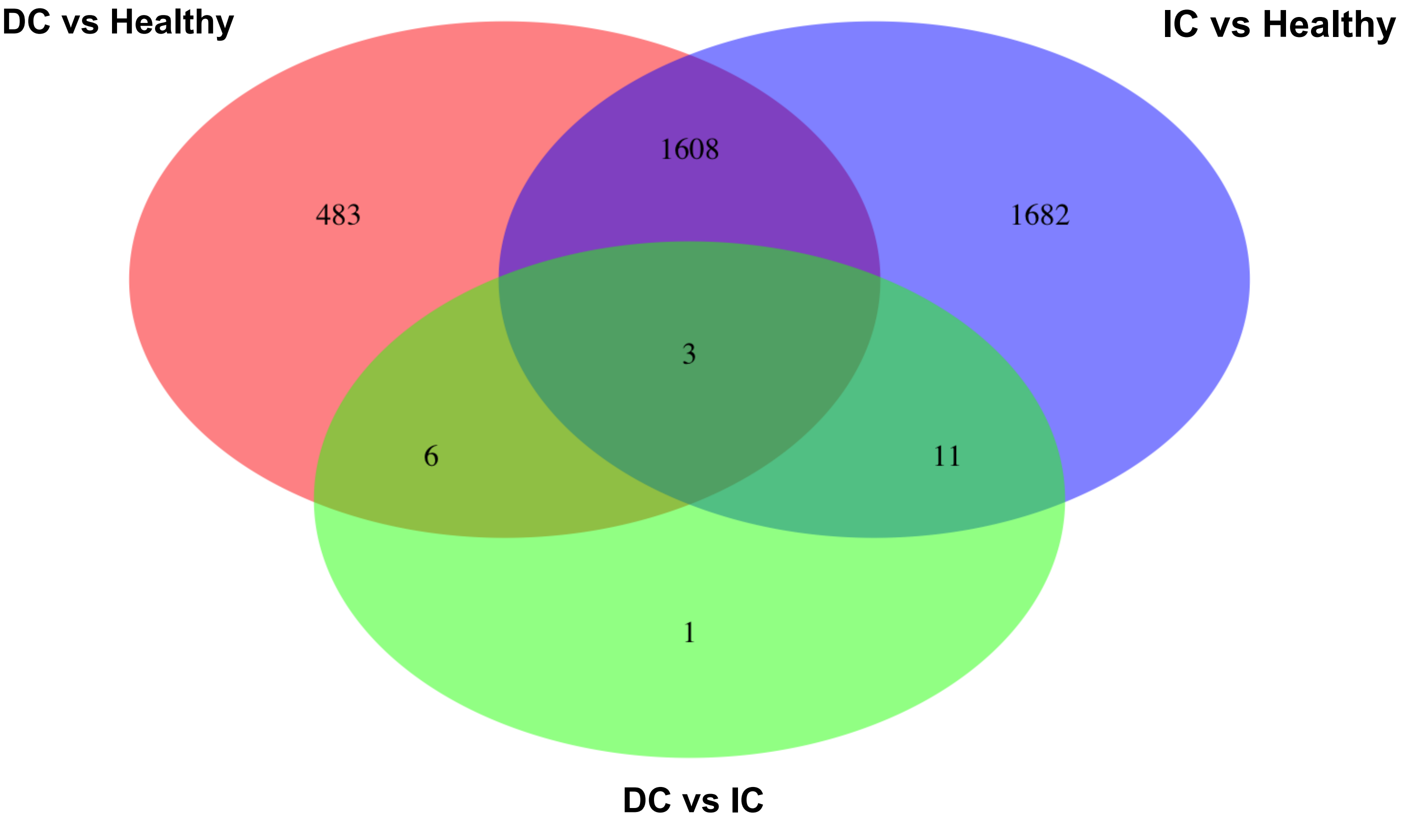

**Supplementary Figure 4.** Venn diagram for significantly changed genes when comparing changed genes across different contrast groups: DC vs Healthy, IC vs Healthy and DC vs IC, where DC - dilated cardiomyopathy and IC - ischemic cardiomyopathy

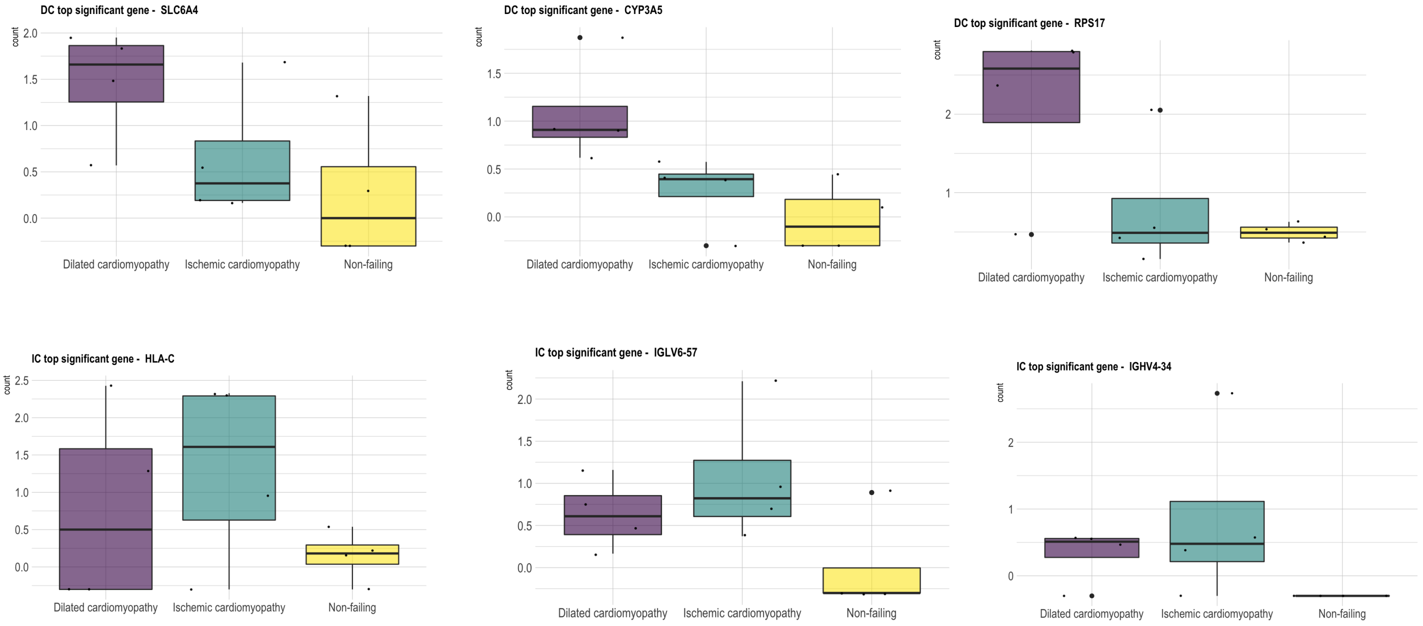


**Supplementary Figure 5.** Log10 scaled gene counts that changed significantly in a specific contrast groups: DC vs Healthy, IC vs Healthy and belonged to the highest log fold change group where DC - dilated cardiomyopathy and IC - ischemic cardiomyopathy. Representative examples shown.

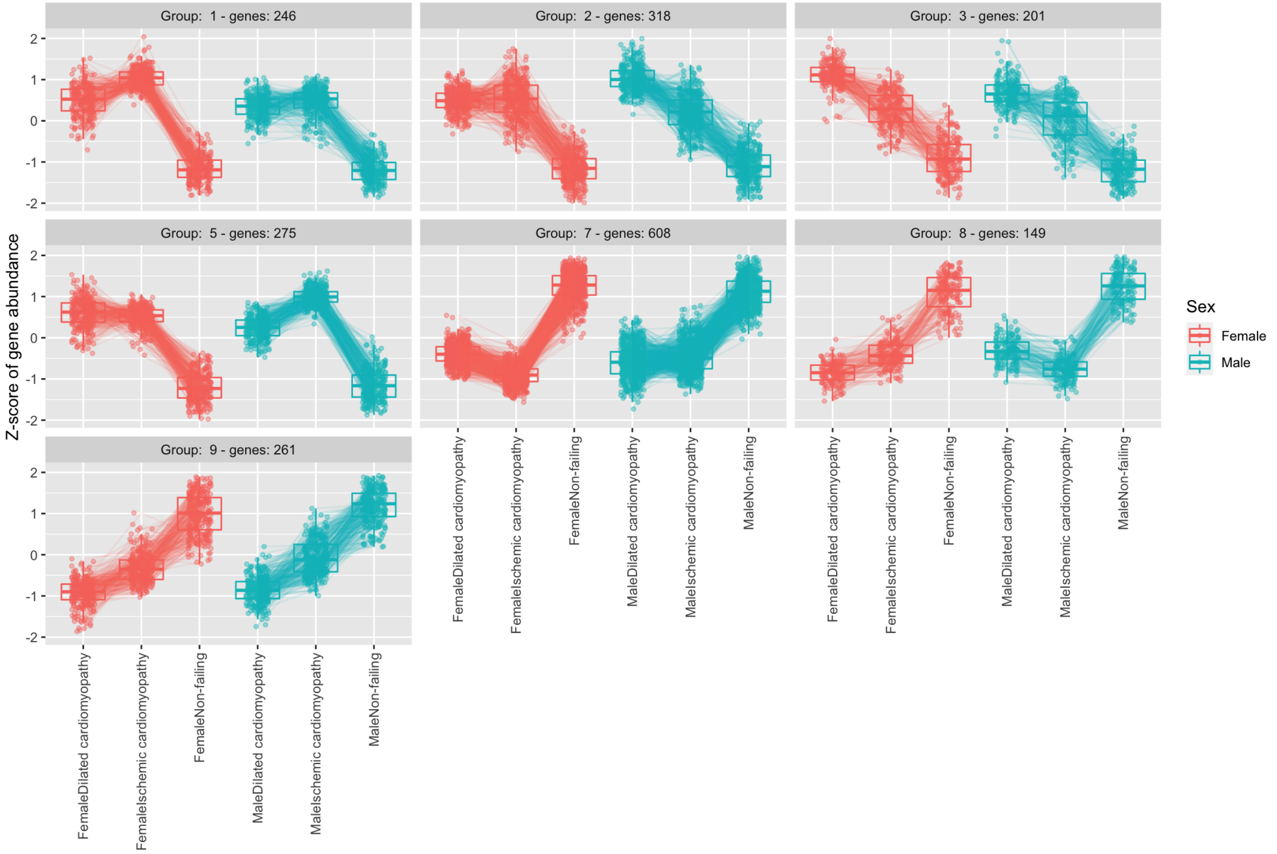


**Supplementary Figure 6.** Scaled gene counts that changed significantly in DC (dilated cardiomyopathy) vs Healthy samples and where expression patterns formed distinct expression clusters across different human heart tissue states when comparing for both genders.

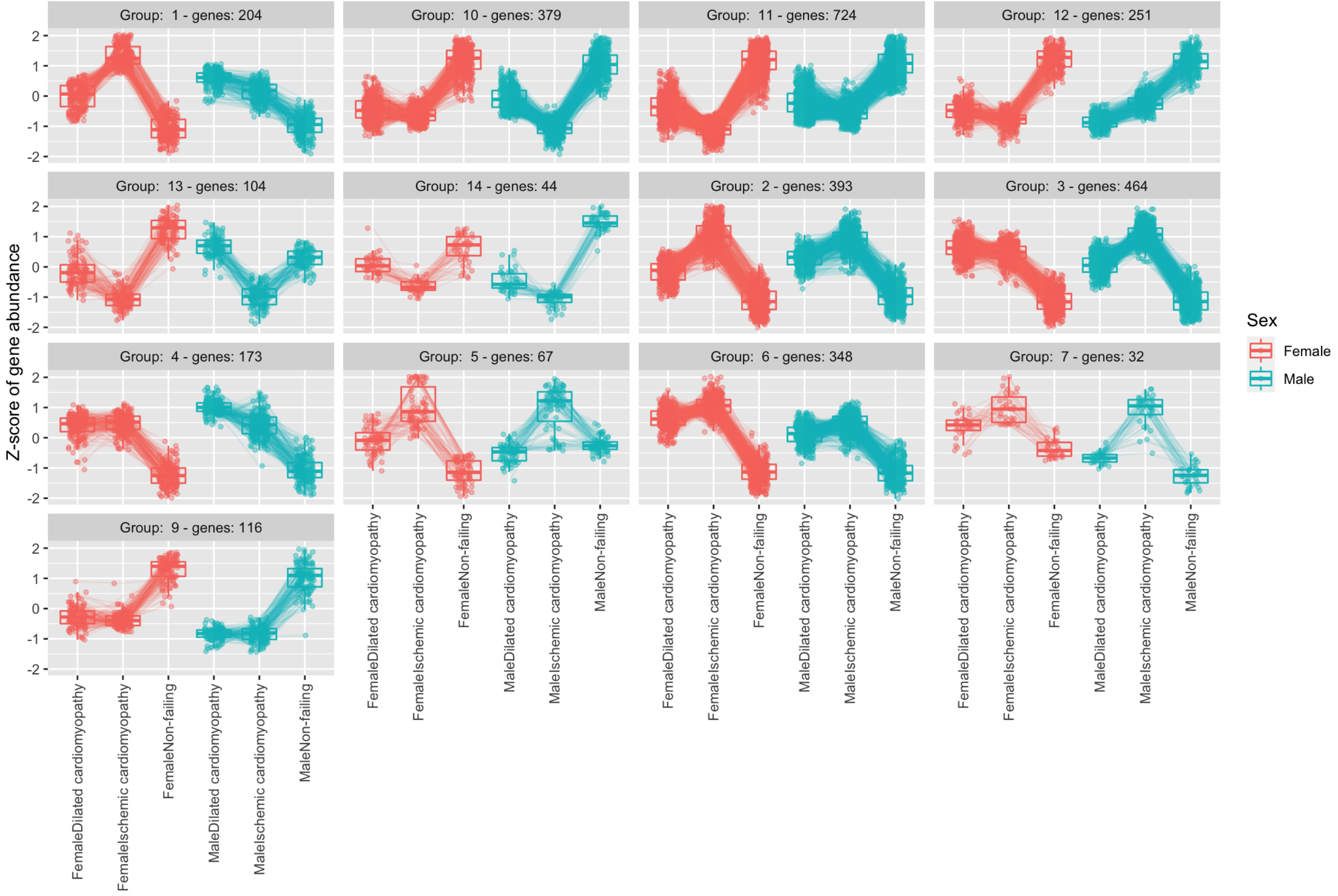

**Supplementary Figure 7.** Scaled gene counts that changed significantly in IC (dilated cardiomyopathy) vs Healthy samples and where expression patterns formed distinct expression clusters across different human heart tissue states when comparing for both genders.

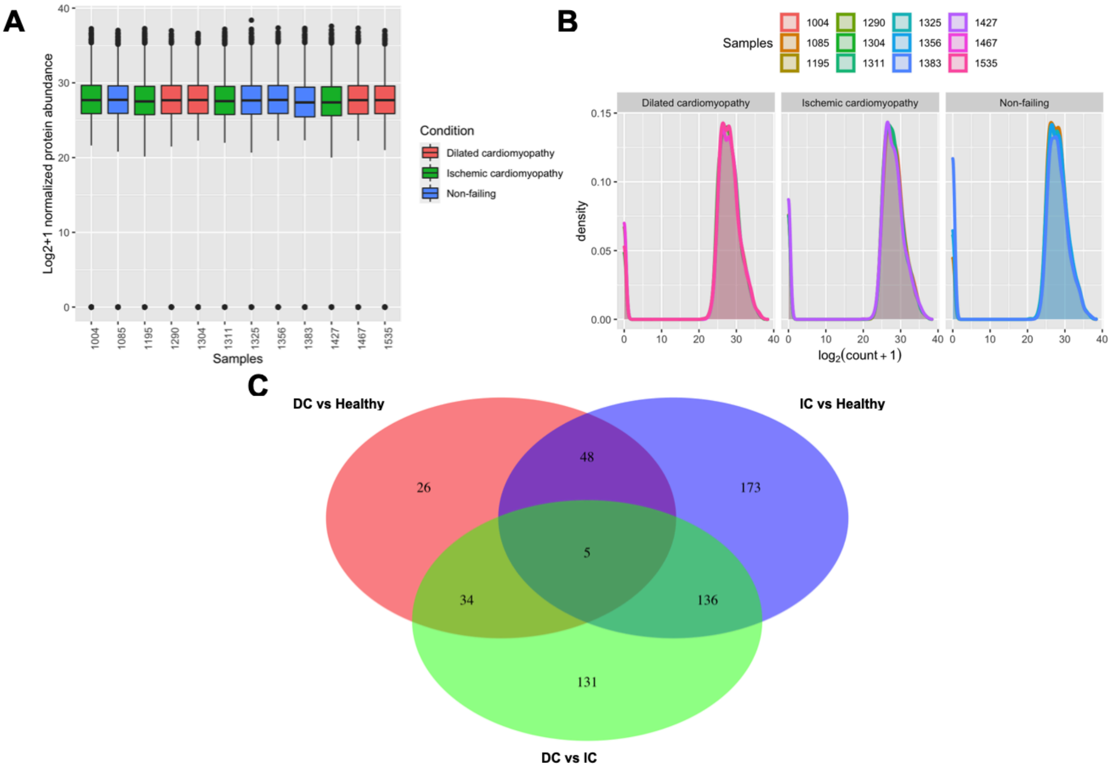


**Supplementary Figure 8.** Protein sample main characteristics: normalised (log2+1 transformed) protein abundance/count (LFQ) values for human left ventricle proteome (PXD008934) (A), density plots of protein samples distribution (B) and shared proteins by different contrast groups are visualised via Venn diagram (C).

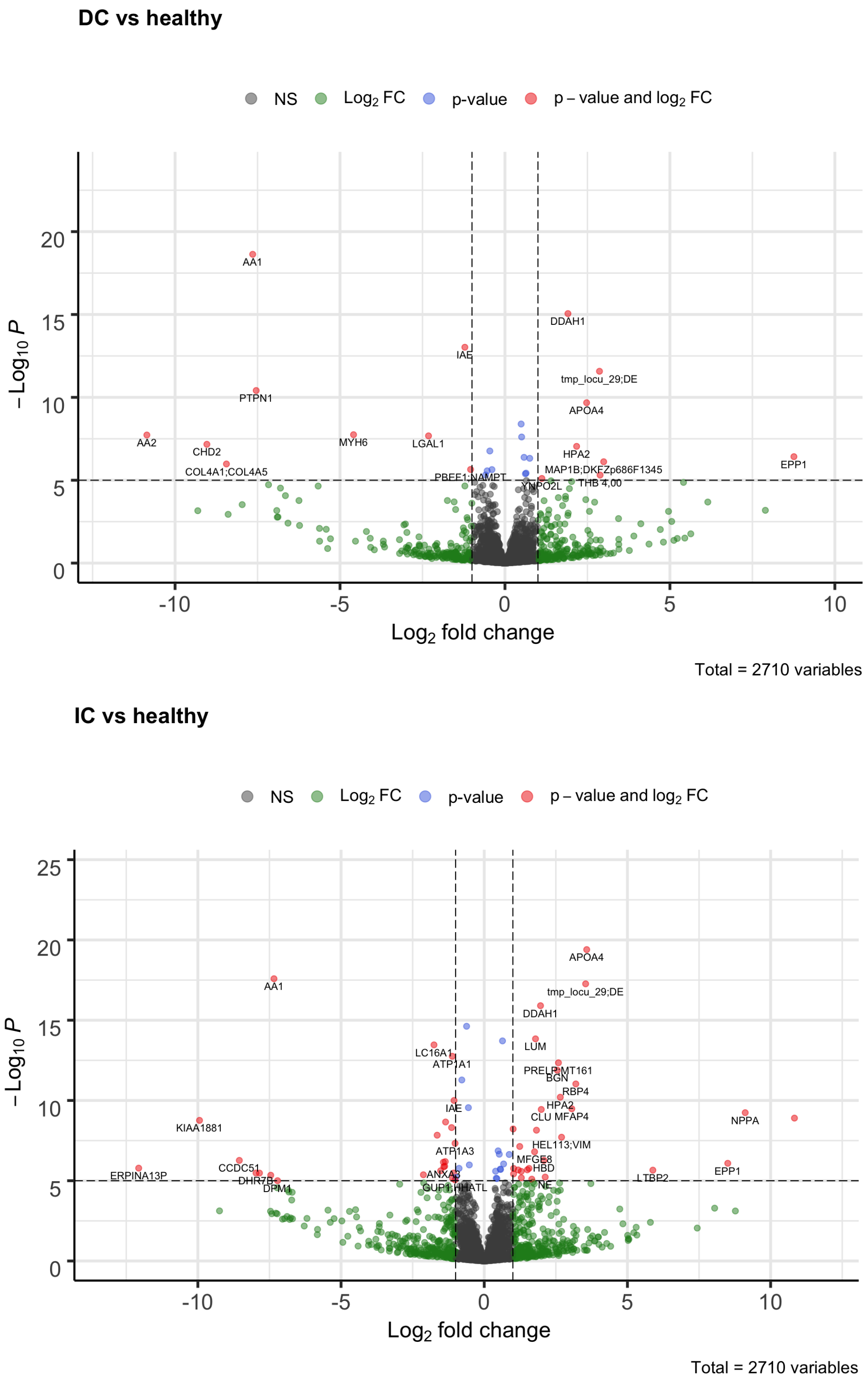


**Supplementary Figure 9.** Volcano plot for human left ventricle proteins (PXD008934) that changed significantly where IC – ischemic cardiomyopathy and DC – dilated cardiomyopathy, FDR p-adjusted<0.00001 and Log fold change (LFC)>|2|.

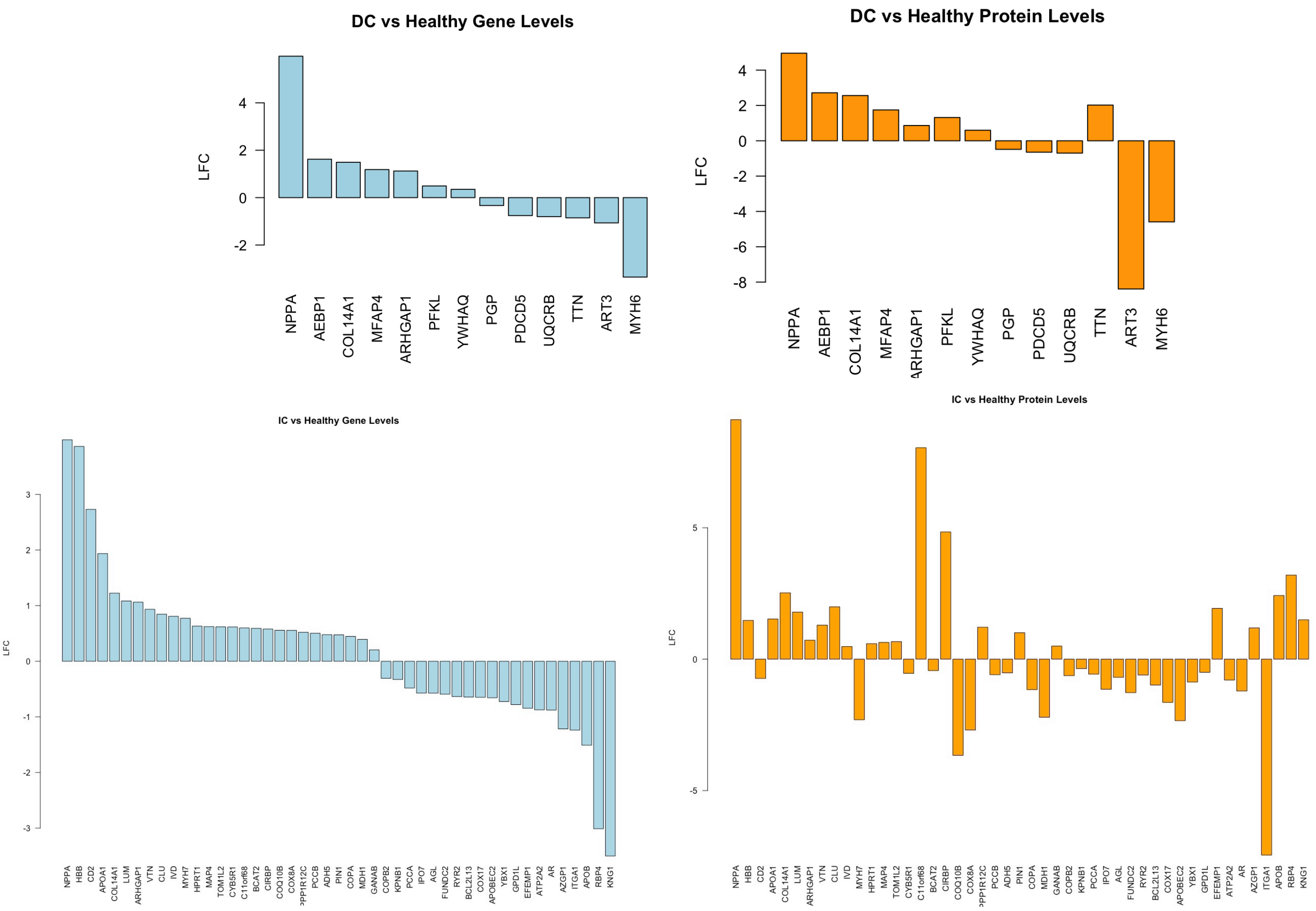


**Supplementary Figure 10.** Gene and protein LFC  values for different contrasts where DC - dilated cardiomyopathy and IC - ischemic cardiomyopathy.

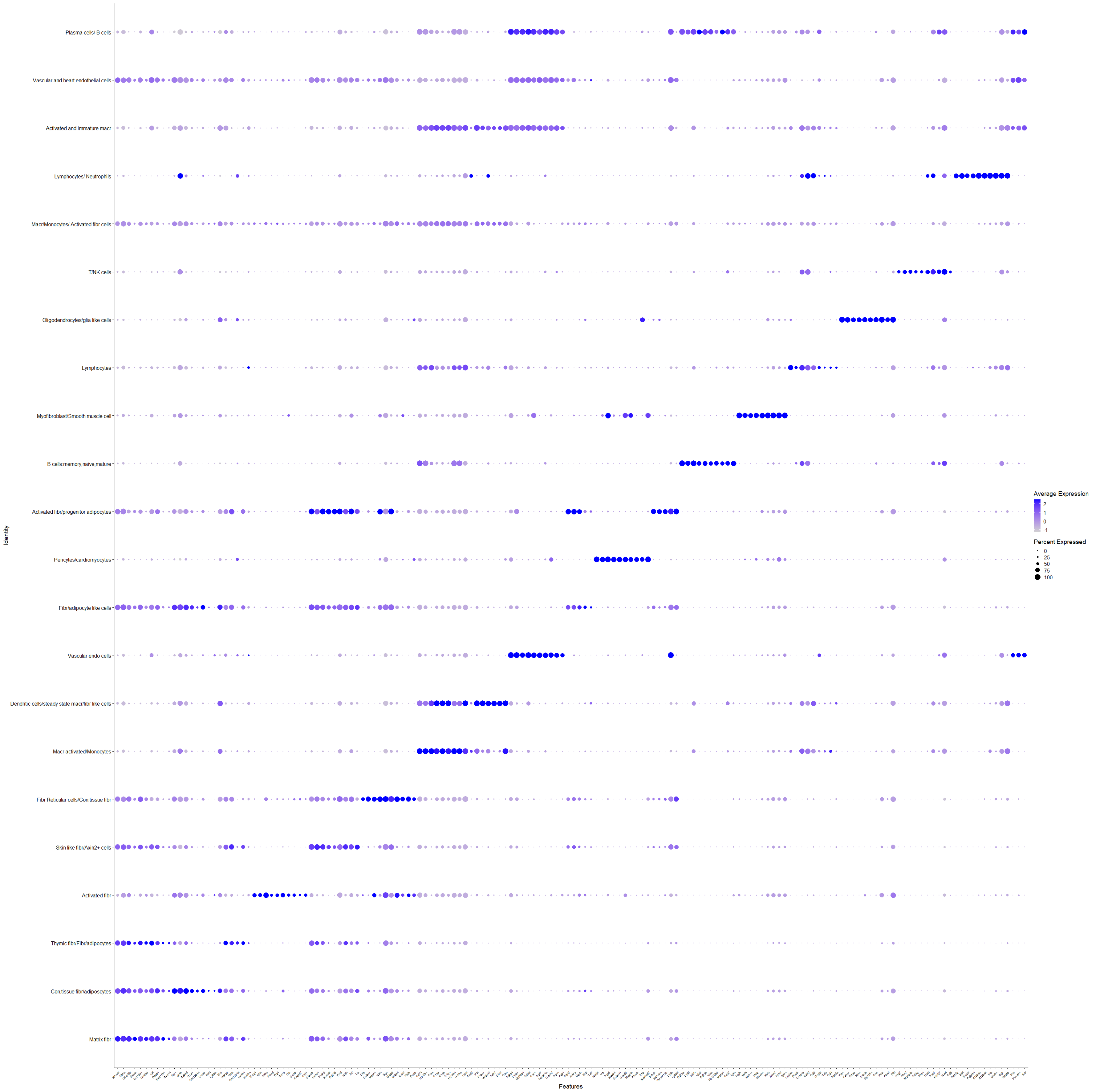


**Supplementary Figure 11.** Mouse non-cardiomyocyte single cell RNA-seq (E-MTAB-6173) cellulome marker gene clusters for the uncovered cell types.

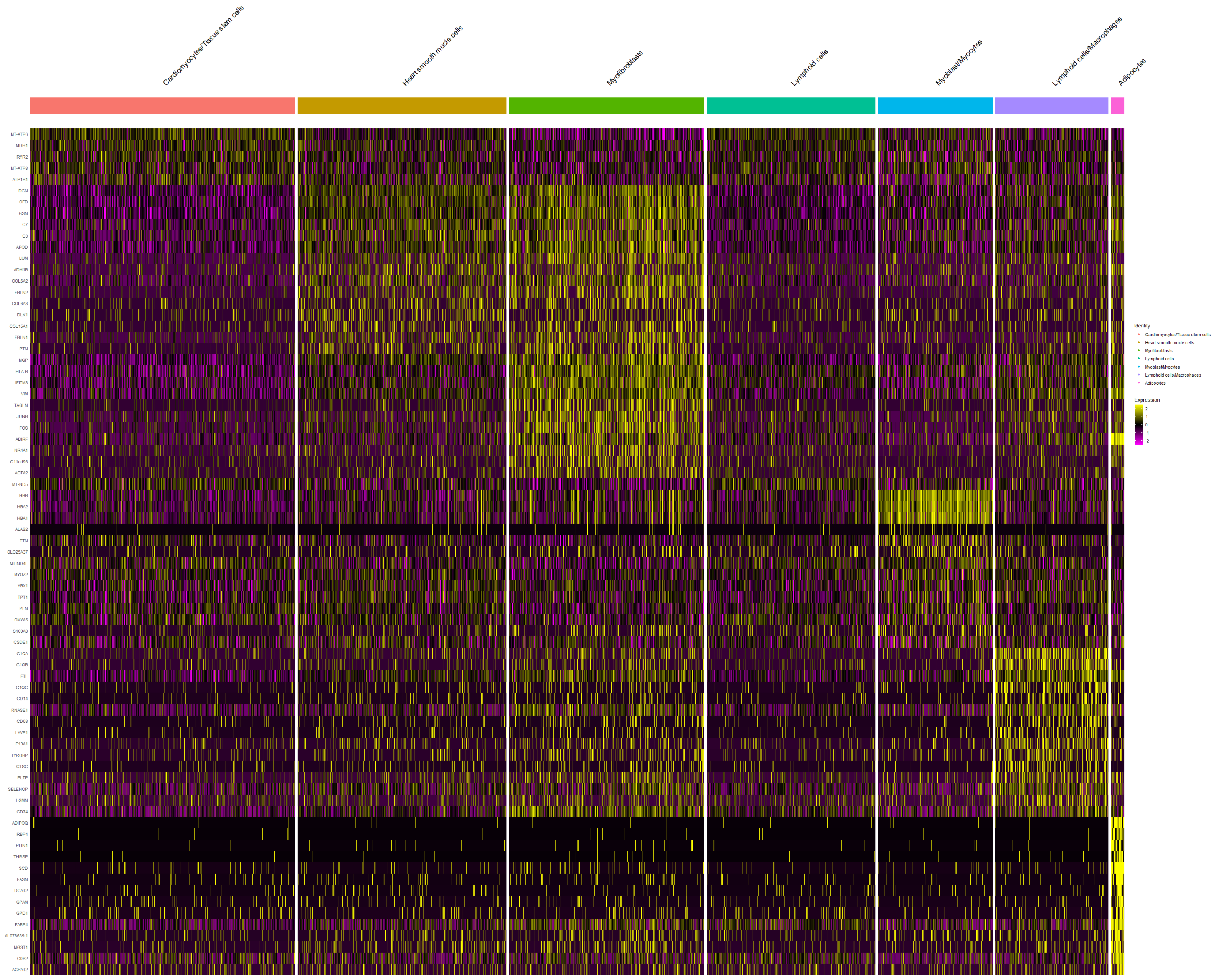


**Supplementary Figure 12.** Human heart left ventricle cellulome marker gene heatmap for the uncovered clusters of different cells.

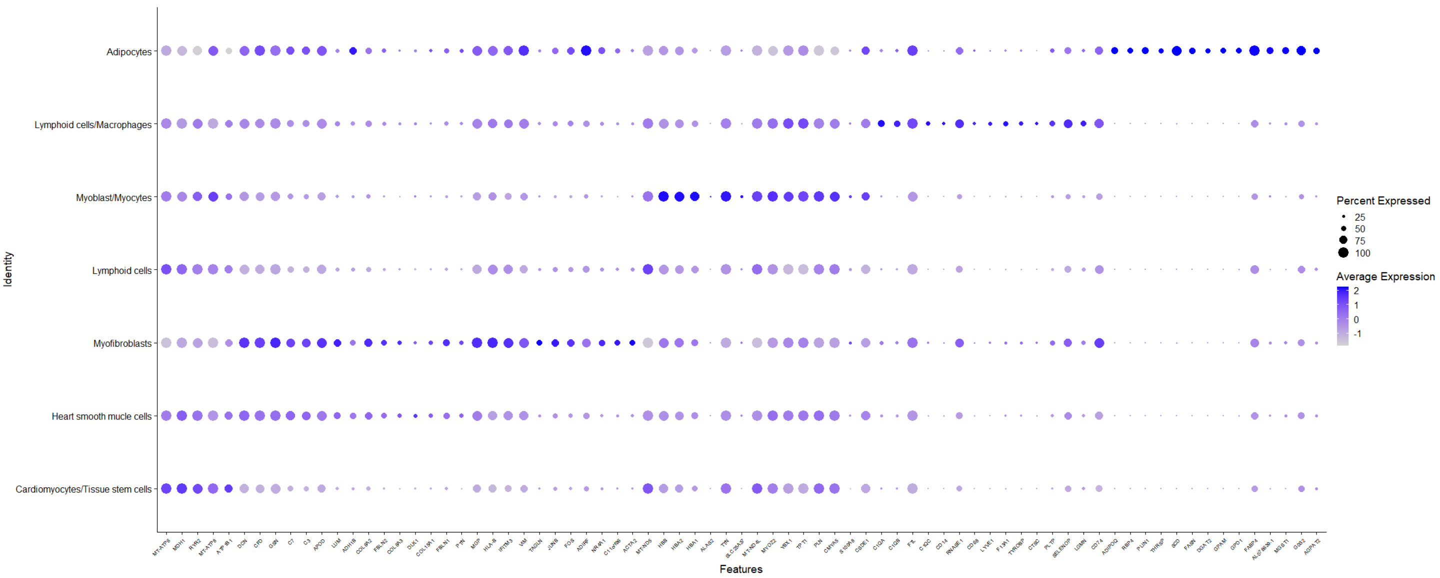


**Supplementary Figure 13.** Human heart left ventricle cellulome marker gene clusters for the uncovered cell types.

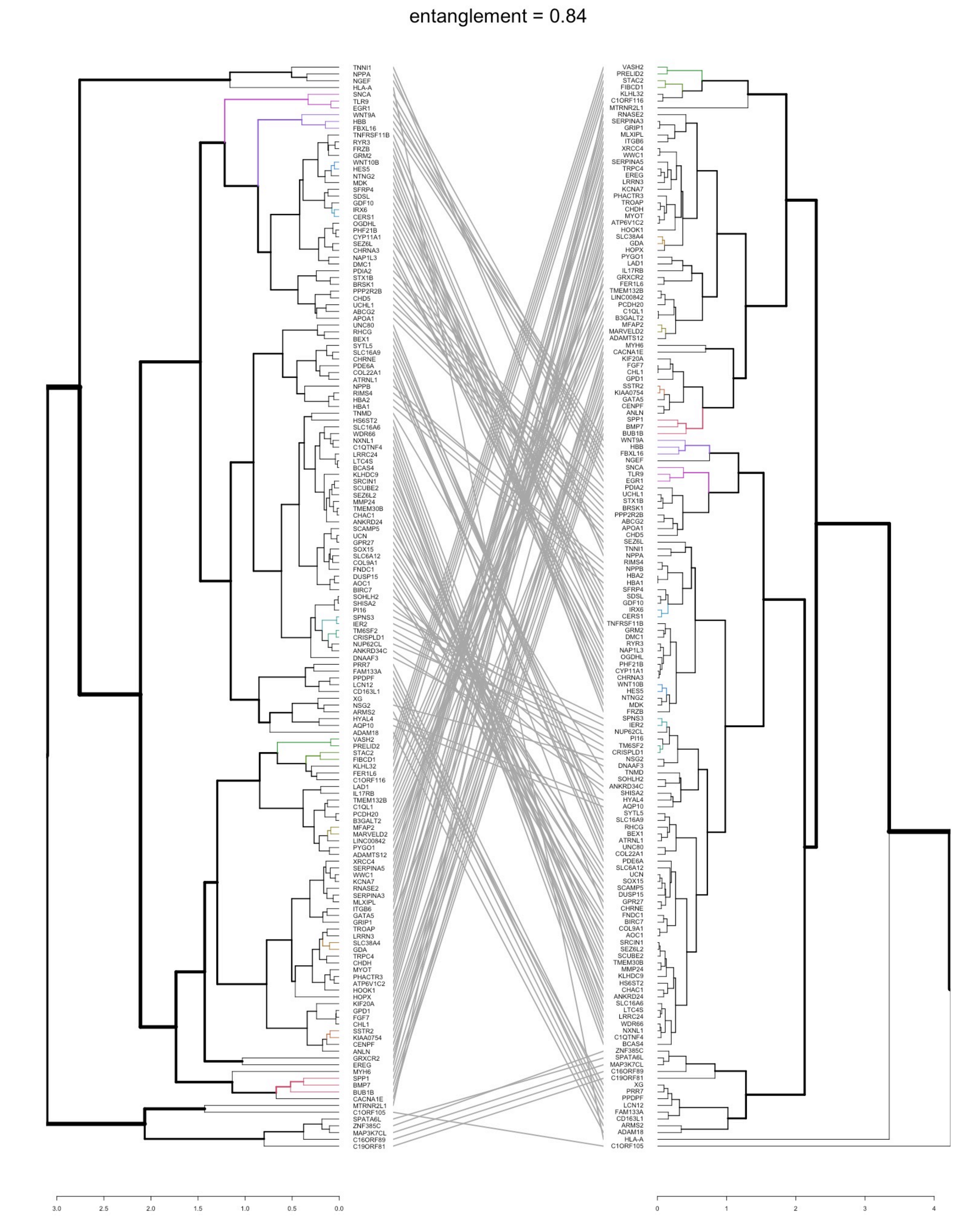
entanglement = 0.84
TNNI1
NPPA
NGEF
HLA-A
SNCA
TLR9
EGR1
WNT9A
HBB
FBXL16
TNFRSF11B
RYR3
FRZB
GRM2
WNT10B
HES5
NTNG2
MDK
SFRP4
SDSL
GDF10
IRX6
CERS1
OGDHL
PHF21B
CYP11A1
SEZ6L
CHRNA3
NAP1L3
DMC1
PDIA2
STX1B
BRSK1
PPP2R2B
CHD5
UCHL1
ABCG2
APOA1
UNC80
RHCG
BEX1
SYTL5
SLC16A9
CHRNE
PDE6A
COL22A1
ATRNL1
NPPB
RIMS4
HBA2
HBA1
TNMD
HS6ST2
SLC16A6
WDR66
NXNL1
C1QTNF4
LRRC24
LTC4S
BCAS4
KLHDC9
SRCIN1
SCUBE2
SEZ6L2
MMP24
TMEM30B
CHAC1
ANKRD24
SCAMP5
UCN
GPR27
SOX15
SLC6A12
COL9A1
FNDC1
DUSP15
AOC1
BIRC7
SOHLH2
SHISA2
PI16
SPNS3
IER2
TM6SF2
CRISPLD1
NUP62CL
ANKRD34C
DNAAF3
PRR7
FAM133A
PPDPF
LCN12
CD163L1
XG
NSG2
ARMS2
HYAL4
AQP10
ADAM18
VASH2
PRELID2
STAC2
FIBCD1
KLHL32
FER1L6
C1ORF116
LAD1
IL17RB
TMEM132B
C1QL1
PCDH20
B3GALT2
MFAP2
MARVELD2
LINC00842
PYGO1
ADAMTS12
XRCC4
SERPINA5
WWC1
KCNA7
RNASE2
SERPINA3
MLXIPL
ITGB6
GATA5
GRIP1
TROAP
LRRN3
SLC38A4
GDA
TRPC4
CHDH
MYOT
PHACTR3
ATP6V1C2
HOOK1
HOPX
KIF20A
GPD1
FGF7
CHL1
SSTR2
KIAA0754
CENPF
ANLN
GRXCR2
EREG
MYH6
SPP1
BMP7
BUB1B
CACNA1E
MTRNR2L1
C1ORF105
SPATA6L
ZNF385C
MAP3K7CL
C16ORF89
C19ORF81
VASH2
PRELID2
STAC2
FIBCD1
KLHL32
C1ORF116
MTRNR2L1
RNASE2
SERPINA3
GRIP1
MLXIPL
ITGB6
XRCC4
WWC1
SERPINA5
TRPC4
EREG
LRRN3
KCNA7
PHACTR3
TROAP
CHDH
MYOT
ATP6V1C2
HOOK1
SLC38A4
GDA
HOPX
PYGO1
LAD1
IL17RB
GRXCR2
FER1L6
TMEM132B
LINC00842
PCDH20
C1QL1
B3GALT2
MFAP2
MARVELD2
ADAMTS12
MYH6
CACNA1E
KIF20A
FGF7
CHL1
GPD1
SSTR2
KIAA0754
GATA5
CENPF
ANLN
SPP1
BMP7
BUB1B
WNT9A
HBB
FBXL16
NGEF
SNCA
TLR9
EGR1
PDIA2
UCHL1
STX1B
BRSK1
PPP2R2B
ABCG2
APOA1
CHD5
SEZ6L
TNNI1
NPPA
RIMS4
NPPB
HBA2
HBA1
SFRP4
SDSL
GDF10
IRX6
CERS1
TNFRSF11B
GRM2
DMC1
RYR3
NAP1L3
OGDHL
PHF21B
CYP11A1
CHRNA3
WNT10B
HES5
NTNG2
MDK
FRZB
SPNS3
IER2
NUP62CL
PI16
TM6SF2
CRISPLD1
NSG2
DNAAF3
TNMD
SOHLH2
ANKRD34C
SHISA2
HYAL4
AQP10
SYTL5
SLC16A9
RHCG
BEX1
ATRNL1
UNC80
COL22A1
PDE6A
SLC6A12
UCN
SOX15
SCAMP5
DUSP15
GPR27
CHRNE
FNDC1
BIRC7
COL9A1
AOC1
SRCIN1
SEZ6L2
SCUBE2
TMEM30B
MMP24
KLHDC9
HS6ST2
CHAC1
ANKRD24
SLC16A6
LTC4S
LRRC24
WDR66
NXNL1
C1QTNF4
BCAS4
ZNF385C
SPATA6L
MAP3K7CL
C16ORF89
C19ORF81
XG
PRR7
PPDPF
LCN12
FAM133A
CD163L1
ARMS2
ADAM18
HLA-A
C1ORF105
3.0
2.5
2.0
1.5
1.0
0.5
0.0
0
1
2
3
4

**Supplementary Figure 14.** Human heart left ventricle bulk RNA-seq shared significantly changed gene (n=160) set between dilated and ischemic cardiomyopathy contrast groups (disease vs healthy state) agglomerative hierarchical clustering based on Log2 Fold Change Score and known interactors returned the shared dendrogram. Coloured branches signify similar clustering patterns.
